# TRPV4 overactivation enhances cellular contractility and drives ocular hypertension in TGFβ2 overexpressing eyes

**DOI:** 10.1101/2024.11.05.622187

**Authors:** Christopher N. Rudzitis, Monika Lakk, Ayushi Singh, Sarah N. Redmon, Denisa Kirdajova, Yun-Ting Tseng, Michael L. De Ieso, W. Daniel Stamer, Samuel Herberg, David Križaj

## Abstract

The risk for developing primary open-angle glaucoma (POAG) correlates with the magnitude of ocular hypertension (OHT) and the concentration of transforming growth factor-β2 (TGFβ2) in the aqueous humor. Effective treatment of POAG requires detailed understanding of interaction between pressure sensing mechanisms in the trabecular meshwork (TM) and biochemical risk factors. Here, we employed molecular, optical, electrophysiological and tonometric strategies to establish the role of TGFβ2 in transcription and functional expression of mechanosensitive channel isoforms alongside studies of TM contractility in biomimetic hydrogels, and intraocular pressure (IOP) regulation in a mouse model of TGFβ2 -induced OHT. TGFβ2 upregulated expression of *TRPV4* and *PIEZO1* transcripts and time-dependently augmented functional TRPV4 activation. TRPV4 agonists induced contractility of TM-seeded hydrogels whereas pharmacological inhibition suppressed TGFβ2-induced hypercontractility and abrogated OHT in eyes overexpressing TGFβ2. *Trpv4*-deficient mice resisted TGFβ2-driven increases in IOP, but nocturnal OHT was not additive to TGFβ-evoked OHT. Our study establishes the fundamental role of TGFβ as a modulator of mechanosensing in nonexcitable cells, identifies the TRPV4 channel as the final common mechanism for TM contractility and circadian and pathological OHT, and offers insights for future treatments that can lower IOP in the sizeable cohort of hypertensive glaucoma patients that resist current treatments.

## Introduction

Primary open-angle glaucoma (POAG), an irreversible blinding disease, afflicts ∼3.5% of the global population (1). Its incidence and severity are proportional to the amplitude and duration of ocular hypertension (OHT) (2, 3), which correlates with retinal ganglion cell dysfunction, neuroinflammation, and oxidative stress (4, 5). Biomechanical factors, glucocorticoids, and the cytokine transforming growth factor-β2 (TGFβ2) contribute to POAG by compromising the funneling of aqueous humor (AH) from the trabecular meshwork (TM) into Schlemm’s canal (SC). Elevated IOP enhances the contractility of the juxtacanalicular TM (JCT), a circumocular tissue comprised of extracellular matrix (ECM) beams populated by mechanosensitive and smooth muscle-like cells- thereby increasing AH outflow resistance. The molecular mechanism that links TM pressure sensing to the contractile response is not known but is likely to underpin the tissue’s sensitivity to compressive, tensile, osmotic, shear and traction forces which collectively regulate expression of numerous TM genes and secretion of ECM proteins (6–12).

The increase in trabecular outflow resistance induced by mechanical stress, glucocorticoids, and TGFβ2 manifests through two distinct components: a dynamic, reversible phase amenable to cytoskeletal and Rho kinase inhibition, and a chronic phase, characterized by transdifferentiation of TM cells into fibrotic and contractile myofibroblasts (16–18). TGFβ2-induced fibrotic remodeling has been linked to POAG: (i) TM cells derived from POAG patients secrete more active TGFβ2 compared to cells isolated from healthy donors (19), (ii) the risk of POAG is proportional to [TGFβ2]_AH_ (20–22), and (iii) ectopic ocular expression of TGFβ2 suffices to induce OHT (23, 24), likely via aberrant secretion of ECM proteins and enhanced TM contractility (25, 26). The cognate TGFβ1 isoform induces similar fibrotic responses in fibroblasts, epithelial, and endothelial cells from heart, kidney, skin, and/or lung, suggesting induction of conserved fibrogrenic programs (27–30). However, the contribution to OHT by ocular TGFβ expression cannot be disambiguated from the changing biomechanical environment: TGFβ release is activated by tissue contractility and tension (31, 32), and TGFβ activity correlates with mechanical stress gradients which may drive a cellular epithelial-mesenchymal transition-like phenotype (EMT; 33-35).

Despite its clinical relevance, our understanding of TM mechanotransduction and its contribution to IOP homeostasis remains rudimentary. Strain and shear stress have been hypothesized to engage primary cilia and integrins, as well as mechanosensitive TRPV4, Piezo1 and TREK-1 channels (36–39), yet it remains unclear whether these mechanosensors regulate TM contractility, are influenced by POAG inducers like TGFβ2 or glucocorticoids, or contribute to chronic fibrosis. Among these, TRPV4 (Transient Receptor Potential Vanilloid isoform 4), a tetrameric channel with P_Ca_/P_Na_ ∼ 10 (40), is strongly expressed in rodent and human TM (36, 51) where it carries the principal component of the pressure-activated transmembrane current and is activated by stretch, shear and swelling (7, 10, 37, 39, 41, 42). Pharmacological inhibition and deletion of the TRPV4 gene alter pressure gradients in the brain, kidney, lung, and bladder (46–50). TRPV4 mutations underpin sensorimotor neuropathies, skeletal dysplasias, retinal degeneration and ocular dysfunction (43–45) while the role of TRPV4 signaling in OHT remains contentious, with evidence suggesting IOP-lowering as well as IOP-elevating effects. TRPV4-dependence of conventional outflow has been linked to diverse downstream effector mechanisms (e.g., eNOS and RhoA activation, phospholipid-cholesterol-caveolin regulation, OCRL inositol-5-phosphatase interaction, modulation of cell-ECM contacts, polyunsaturated fatty acid release, and Piezo1 signaling; 7, 37, 41, 52–55) and thus leads towards testable hypotheses: if TM-intrinsic TRPV4 sustains steady-state normotension, promotes outflow via eNOS-dependent TM relaxation and mitigates TGFβ2 -driven fibrosis (7, 52), TRPV4 inhibition should induce OHT. Conversely, if TRPV4 exacerbates OHT, its blockade and deletion should reduce IOP.

In this study, we tested these hypotheses through investigation of reciprocal TRPV4- TGFβ2 interactions that perpetuate the vicious feedback loop between mechanical stressors, TM contractility, and OHT. We demonstrate that inhibition and deletion of TRPV4 lower IOP in TGFβ2 overexpression-induced and circadian OHT models and suppress TM contractility in TGFβ2-treated biomimetic hydrogels. The cytokine promoted upregulation of EMT-associated genes alongside increased transcription and activity of TRPV4, potentially sensitizing TM cells to physiological mechanical cues. Although TRPV4 activity was required to maintain OHT under physiological (nocturnal) and pathological (cytokine-induced) conditions, their respective IOP elevations were not additive, suggesting convergence on a shared common pathway. Collectively, these findings position TRPV4 as a critical nexus of TGFβ2 -induced TM contractility and IOP dysregulation. As such, TRPV4 perpetuates the vicious feedback loop between mechanical stressors and TM contractility and thus represents an ideal therapeutic target in glaucoma cases that resist current treatments.

## Results

### TGFβ2 drives overexpression of genes that encode fibrotic markers and mechanosensitive ion channels

Human TM cells respond to TGFβ2 with increased biosynthesis, deposition and degradation of ECM, altered autophagy, upregulation of F-actin stress fibers, a-smooth muscle actin (aSMA) (19, 25, 26, 56, 57), but it is unclear whether cells undergoing TGFβ2-induced fibrotic remodeling also exhibit altered capacity for sensing and transduction of mechanical stimuli. We profiled genes that encode known TM mechanochannels together with a selection of key cytoskeletal, ECM, and fibrotic markers in primary TM cells (pTM) isolated from 3-7 donors without history of visual dysfunction (Figure 1*A*-*C*). Five-day exposure of pTM cells to a physiological concentration of TGFβ2 (1 ng/mL) increased the expression of EMT-promoting transcription factor SNAI1 (*SNAIL1*, *P* = 0.0094) and fibronectin (*FN1, P* = 0.0263), while expression of connective tissue growth factor *2* (*CCN2*, alternatively *CTGF*) was elevated in 5/5 pTM cell strains without reaching significance (*P* = 0.0909). Expression of fibroblast-specific protein 1 (*FSP1*, a calcium-binding fibroblast marker), yes-associated protein 1 (*YAP1*, a stiffness induced hippo-pathway transcription factor) and *ACTA2* (αSMA, associated with cell contractility) was not consistently impacted by TGFβ2 while transcription of myocilin (*MYOC*) decreased across 4/4 pTM strains (*P* = 0.0055) (Figure 1*B*). Indicative of feedback inhibition (58), TGFβ2-treatment downregulated transcript levels of transforming growth factor beta receptor 2 (*TGFBR2*, *P* = 0.0219) and upregulated expression of autoinhibitory SMAD family protein 7 (*SMAD7*, *P* = 0.0461) without affecting *SMAD2* or *SMAD3* expression. TGFβ2 thus promotes selective upregulation of ECM and fibrosis-related genes together with cell dedifferentiation and activation of autoregulatory SMAD mechanisms.

**Figure 1:**
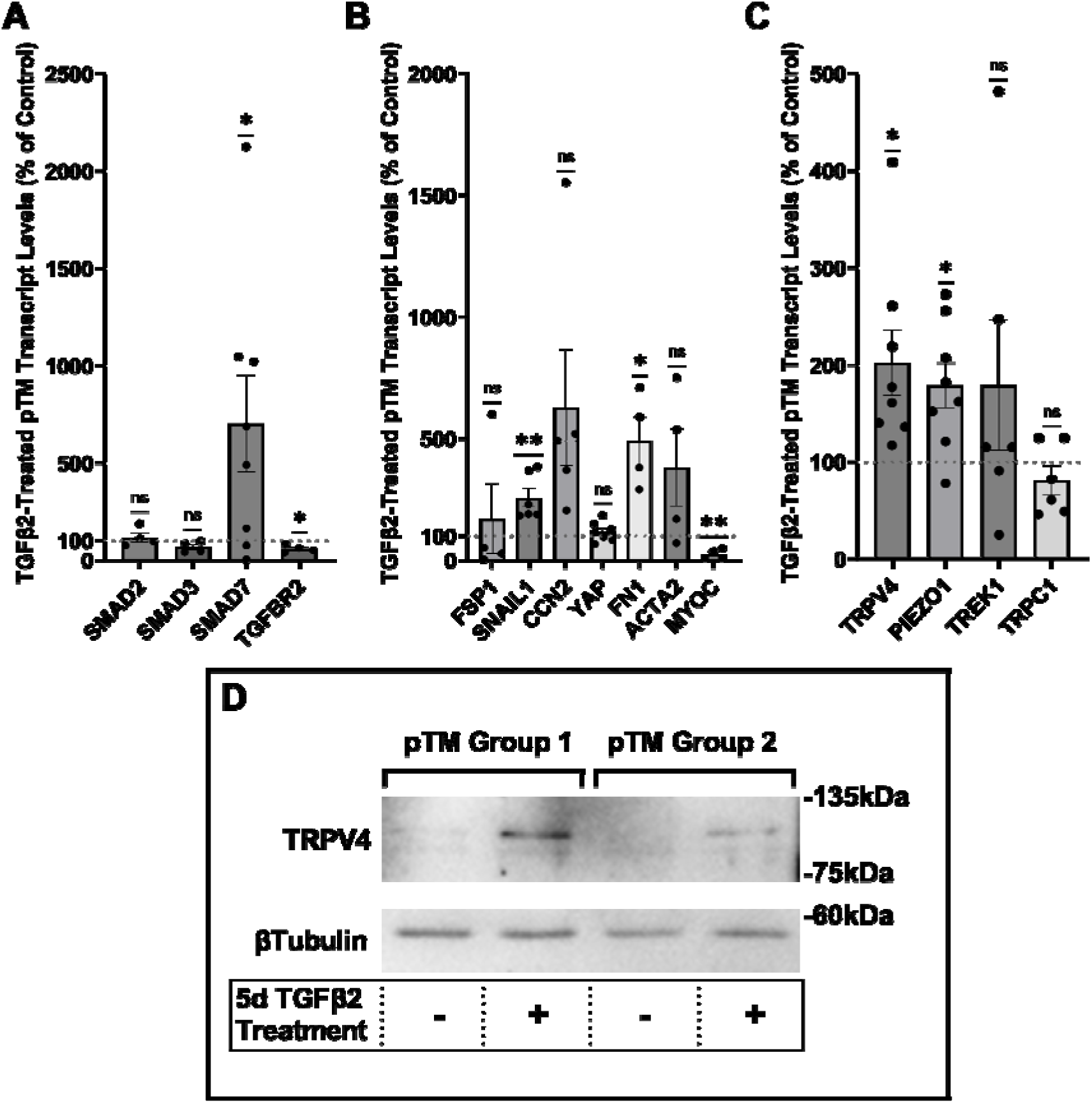
TGFβ2 induces a fibrotic phenotype in pTM cells and increases expression and membrane insertion of the TRPV4 channel. (**A**-**B**) Five-day TGFβ2 treatment (1ng/mL) significantly altered expression of TGFβ pathway effectors, cytoskeletal machinery, and canonical fibrotic markers. (**C**) TGFβ2 treatment significantly increased *TRPV4* and *PIEZO1* expression, but not *TREK1* and *TRPC1* expression. Mean ± SEM shown. N = 4 - 8 experiments, each gene tested in 3-7 different pTM strains (See Table 1). Two-tailed one sample t-test of TGFβ2-induced gene expression levels as a percent of control samples. (**D**) Isolation of membrane proteins from two separate pooled pTM samples suggests TGFβ2 treatment drives increased TRPV4 membrane insertion. N = 2 independent pooled samples, 3 pTM strains were pooled per sample. *\* P* < 0.05, *\*\* P* < 0.01.

**Table 1:**
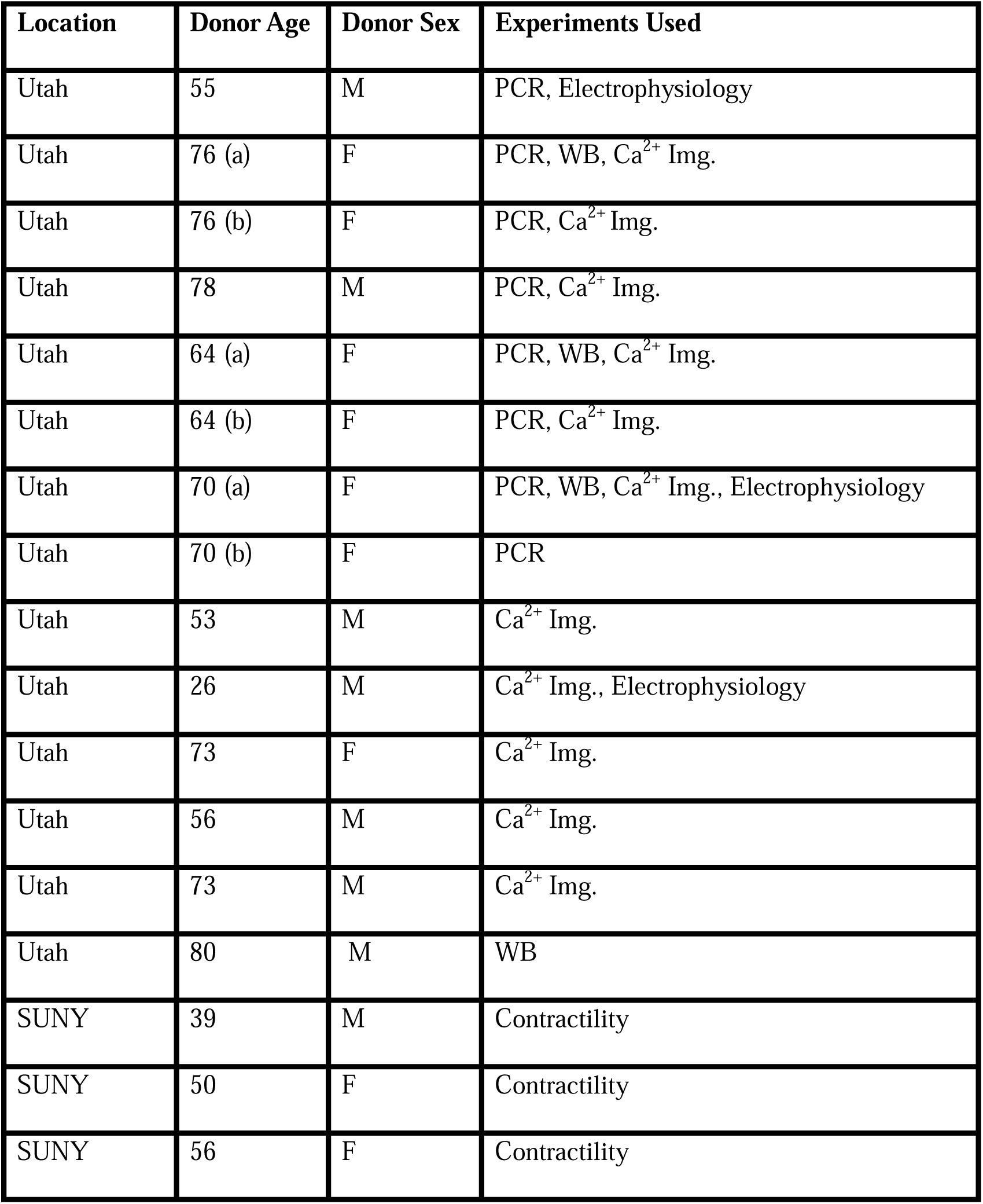
Donor information for primary human trabecular meshwork (pTM) strains used in this study.

Analysis of genes encoding mechanosensitive channels implicated in outflow modulation (36, 39, 59, 60) showed a 102.5% increase in expression of *TRPV4* (*P* = 0.0193) and 78.9% increase in *PIEZO1* expression (*P* = 0.0114) across 8 replicates including 7 distinct pTM strains. (Figure 1*C*). Conversely, TGFβ2 exposure did not affect expression of the *TRPC1* gene (*P* = 0.261) and had variable, strain-dependent effects on transcript levels *of KCNK2 (P* = 0.293, encoding the TREK-1 channel). Thus, TGFβ2 promotes selective transcriptional upregulation of genes that encode a subset of mechanosensitive proteins alongside fibrotic upregulation and cell dedifferentiation. Finally, we tested whether TGFβ2-induced upregulation of TRPV4 and Piezo1 is TRPV4-dependent, however, inclusion of the selective TRPV4 inhibitor HC067-47 (HC-06; 5 mM) had no effect on transcriptional upregulation compared to TGFβ2 treatment alone (SI Appendix Figure S1).

### TGFβ2 exposure time-dependently augments TRPV4-mediated current and Δ[Ca^2+^]_i_

To assess the functional relevance of TGFβ2-dependent transcriptional upregulation we determined the membrane expression and functional activation of TRPV4, which mediates the pressure-activated current and calcium signaling, regulates cytoskeletal dynamics and modulates conventional outflow resistance in vitro (37, 41). TGFβ2 exposure produces an increase in levels of membrane-bound TRPV4 protein (Figure 1*D*) in western blot of two grouped pTM membrane protein samples. While low amounts of TRPV4 were visible in the membrane fractions in control samples, TGFβ2 treatment produced an increase in the higher weight TRPV4 band, suggesting there may be isoform-specific TGFβ2-induced responses and increased TRPV4 translation leading to elevated TRPV4 trafficking, membrane insertion and/or lipid raft interaction (52).

Functional expression was assessed by tracking [Ca^2+^]_i_ changes in cells exposed to the selective TRPV4 agonist GSK1016790A (GSK101, 10 nM) using ratiometric Fura2-AM Ca^2+^ dye, with TGFβ2-treated and control cells tested on the same day. All pTM strains responded to GSK101 with robust [Ca^2+^]_i_ increases which reached peak within 5 min before the majority of responding cells gradually decreased to a steady plateau (Figure 2*C*). TGFβ2-treated cells exhibited a remarkable potentiation of GSK101-evoked [Ca^2+^]_i_ responses compared to control cells, with 5/5 cell strains showing an increase in the Δpeak/baseline F_340_/F_380_ response equivalent to 258.4% ± 61.7% of the control response in (*P* = 0.0046) (Figure 2*A-B*). The fraction of GSK101 responders and the overall time course of responses between groups were not significantly different, indicating that TRPV4 potentiation primarily affects TRPV4-expressing cells. Thus, TGFβ2 treatment promotes TRPV4 expression and functional activity.

**Figure 2:**
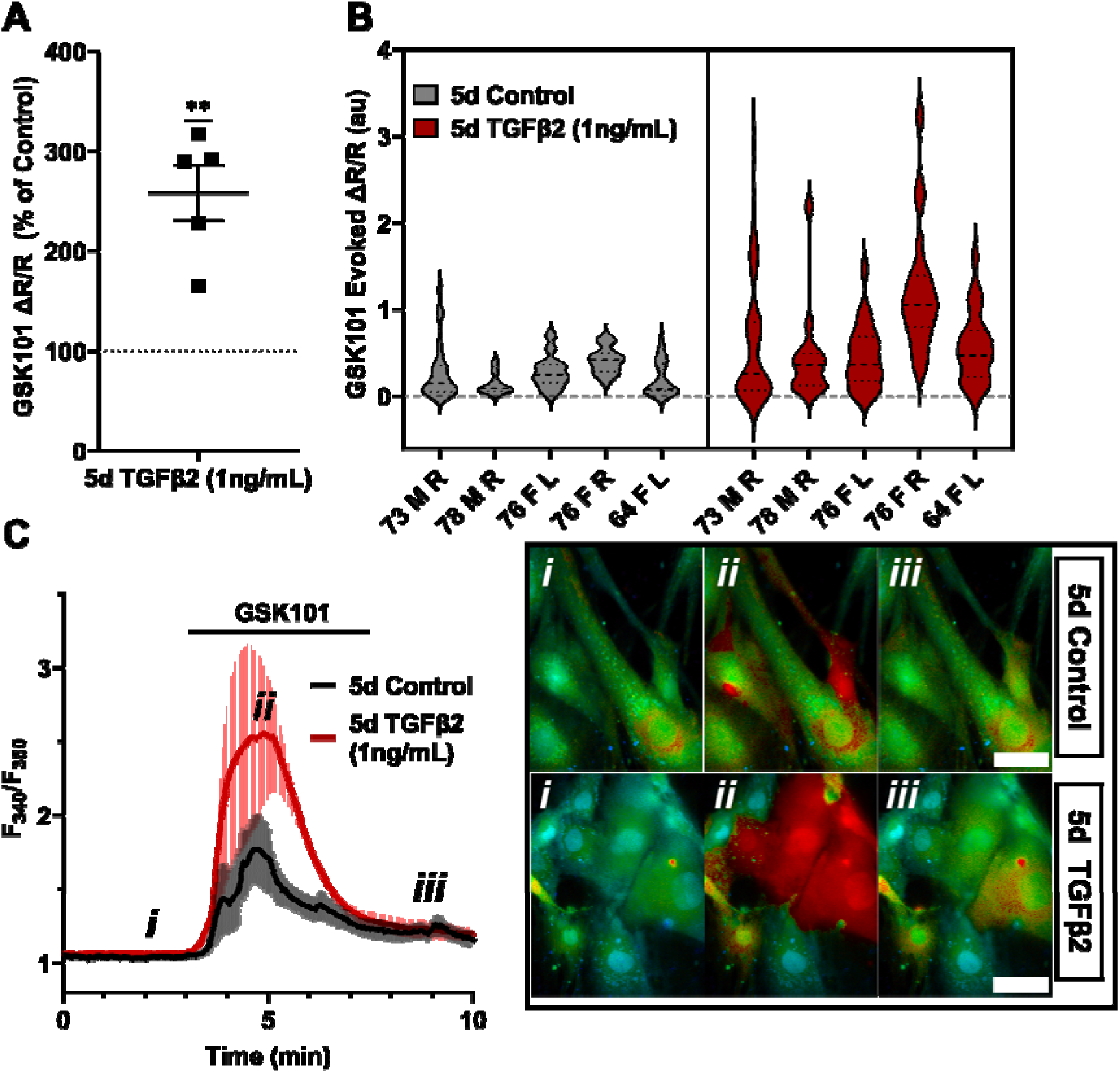
TRPV4-mediated Ca^2+^ influx is potentiated by five-day TGFβ2 treatment. (**A**) Five-day TGFβ2 treatment (1 ng/mL) increased TRPV4 agonist-induced (GSK101, 10 nM) Ca^2+^ influx in pTM cells compared to serum-free media alone treated cells tested on the same day (N = 5 pTM strains, n = 3 - 5 slides/condition/day, individual data points over mean ± SEM). Two-tailed one sample t-test of TGFβ2-treated cell average GSK101 response as a percent of control samples from the same pTM strain on the same day. (**B)** Violin plots showing the distribution of GSK101-induced Ca^2+^ responses for each pTM strain tested in A. Thick dashed line indicates mean, while light dashed line indicates quartiles. (**C**) Representative traces showing TRPV4 agonist-induced Ca^2+^ influx (seen as an increase in F_340_/F_380_) in pTM (mean ± SEM of 4 representative cells/ group), alongside example Fura-2-loaded pTM cells before (i), during (ii), and after (iii) GSK101 application. Scale bar = 50 µm. *\*\* P* < 0.01

To gain insight into the time- and dose-dependence of TGFβ2-dependent TRPV4 signaling modulation pTM cells were treated for 24 hours, at 1 ng/mL and 5 ng/mL concentrations of TGFβ2. GSK101-stimulated Ca^2+^ influx was not significantly increased by 24-hour TGFβ2 treatment at 1 ng/mL (Δpeak/baseline F^340^/F^380^ = 117.0% ± 23.6% of control) or 5 ng/mL (Δpeak/baseline F^340^/F^380^ = 133.6% ± 34.5% of control) (Figure 3; SI Appendix Figure S2); the potentiation of both was significantly lower relative to the five-day 1 ng/mL TGFβ2 treatment (*P* < 0.0011; Figure 3*A*). GSK101 evoked a moderately outwardly rectifying nonselective current (I_GSK_-I_baseline_) with reversal potential at ∼0 mV (Figure 3*C*). While its amplitude was variable, mean current density consistently increased in cells treated for 1 day with TGFβ2 (n = 10; 5 ng/mL) relative to the control group (n = 11). The potentiating effect of TGFβ2 on TRPV4 activity thus appears to be time-dependent but is significant after chronic exposure to relatively low-dose TGFβ2.

**Figure 3:**
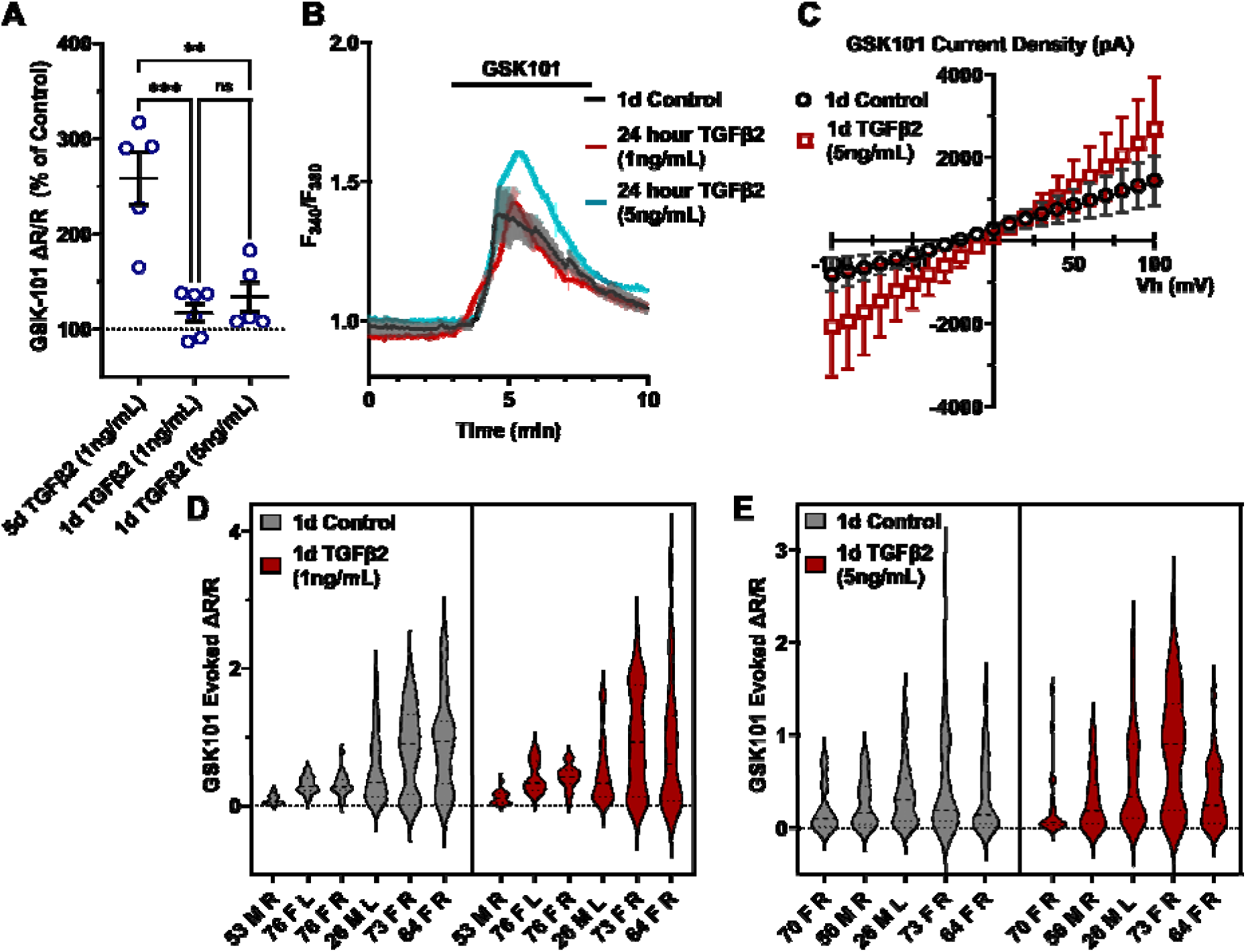
TGFΒ2-induced TRPV4 potentiation is not seen at a shorter time period, regardless of treatment strength. (**A**) TGFβ2 treatments for 24 hours at 1 ng/mL (N = 6 pTM strains, n = 3 - 5 slides/condition/day) or 5 ng/mL (N = 5 pTM strains, n = 3 - 5 slides/condition/day) did not show potentiation of GSK101-evoked TRPV4 Ca^2+^ influx (SI Appendix Figure S2) and were significantly lower than cells treated with TGFβ2 for 5d at 1ng/mL (5d TGFβ2 results from Figure 2A). Individual data points over mean ± SEM. One-way ANOVA with Tukey’s multiple comparisons test, statistics for individual 1d treatment groups compared to control groups shown in Figure S1. (**B**) Representative traces for GSK101 response following 24-hour TGFβ2 treatment, traces show mean ± SEM of 3-4 cells. (**C**) Average current density in response to GSK101 (24-hour control: n = 11 cells, 24-hour TGFβ2: n=10 cells) shows generally increased current in TGFβ2-treated cells. Data shows mean ± SEM (**D - E**) Violin plots of individual cell strains shown in **A**. Thick dashed line indicates mean, while light dashed line indicates quartiles. *\*\* P <* 0.01, *\*\*\* P <* 0.001

### TGFβ2-Induced TM contractility requires TRPV4 activation

The IOP-lowering effectiveness of Rho kinase inhibitors and latrunculins (57, 61–63) indicates that sustained increases in outflow resistance require tonic actin polymerization and contractility. TGFβ2 drives the TM myofibroblast contractile response (57) while the role of mechanosensation remains unknown. To ascertain whether TRPV4 upregulation (Figures 1-2) contributes to the contractile response, we seeded pTM cells into high-compliance Type I collagen hydrogels (57) (Figure 4, SI Appendix Figure S3-4). Hydrogels that were incubated with TGFβ2 (5 ng/mL) showed profound increases (*P* < 0.0003) in the rate and the magnitude of contraction at all time points (Figure 4, SI Appendix Figure S3-4). Simultaneous treatment with HC-06 (5 µM) significantly reduced the extent of TGFβ2-induced TM contractility (*P* < 0.0001). To determine whether TRPV4 activation is sufficient to induce the contractile response, the antagonist was washed out and hydrogels supplemented with GSK101 (25 nM). 15 minutes post-treatment, the constructs responded to the agonist with transient contraction (Figure 4*C*; SI Appendix Figure S3, *P* < 0.01), with a time course mirroring GSK101-induced [Ca^2+^]_i_ elevations (Figures 2-3). The effects of TRPV4 inhibition and activation were consistent across all pTM strains tested (N = 3). TRPV4-mediated Ca2^2+^ influx is therefore sufficient to induce TM contractility and necessary for pTM hypercontractility induced by TGFβ2.

**Figure 4:**
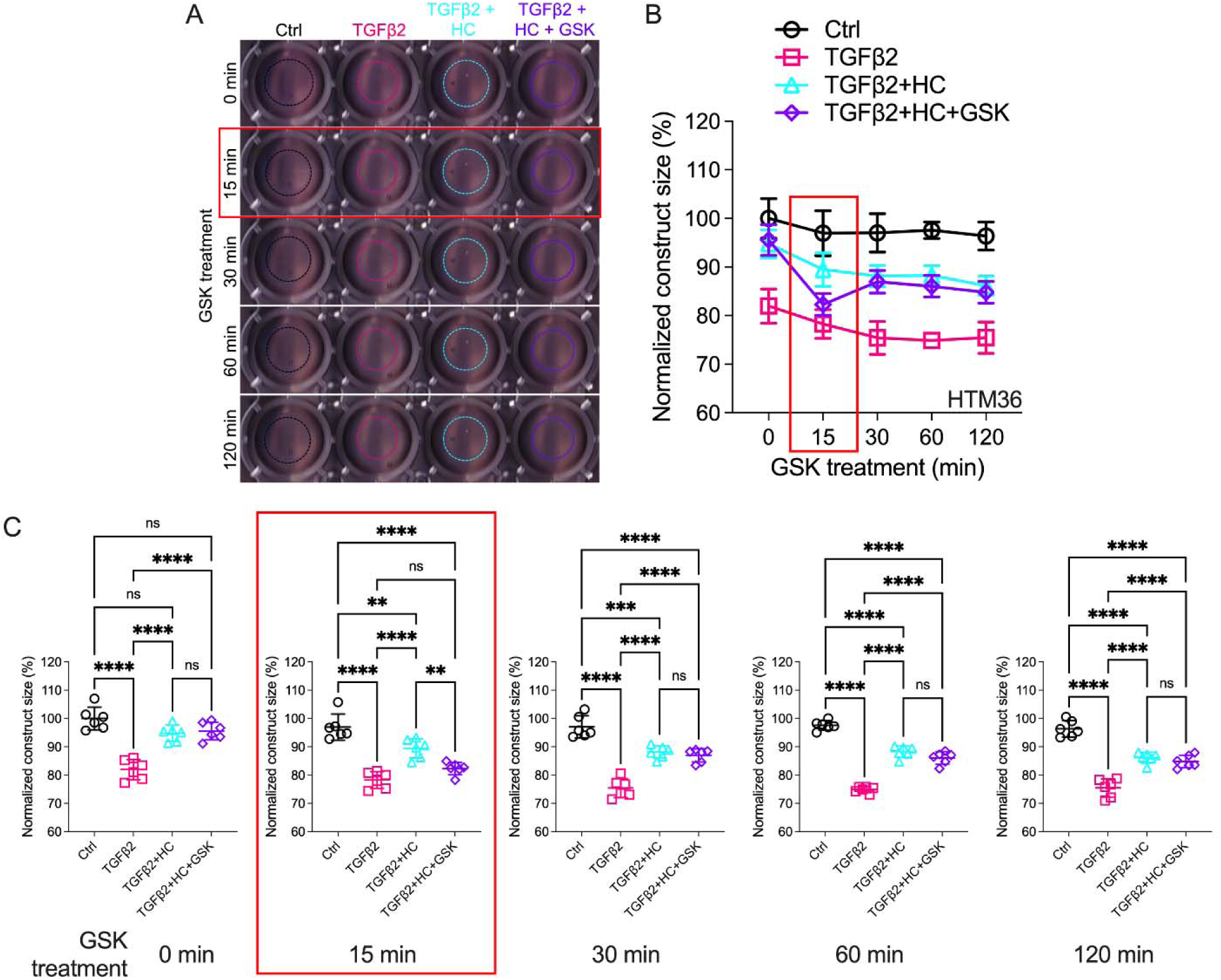
Effects of TRPV4 inhibition/activation on TGFβ2-induced contraction of TM cells. (**A**) Representative longitudinal 24-well plate scans of collagen type I hydrogels seeded with pTM subjected to the different treatments (dashed lines outline size of contracted constructs). (**B**) Longitudinal quantification of hydrogel construct size compared to the control group at the 0 minute time point. (**C**) Detailed comparisons between groups at each experimental time point. n = 6 hydrogels/group. One-way ANOVA with Tukey’s multiple comparisons test, data in (**B & D**) shows individual data points over mean ± S.D. One pTM strain shown: TGFβ2-induced contractility induction, HC-06-mediated rescue of hypercontractility, and GSK101-induced transient (15 min) contraction were consistent across (3/3) pTM strains tested (SI Appendix Figure S3). *\*\* P* < 0.01, *\*\*\* P* < 0.001, *\*\*\*\* P* < 0.0001.

### TRPV4 activation is required to maintain TGFβ2-induced OHT

To test whether TRPV4 contributes to TGFβ2 induced ocular hypertension (OHT) *in vivo*, we utilized the lentiviral TGFβ2 overexpression model developed by Patil et al. (23). Adult C57BL/6J mice (N = 5) were intravitreally injected with lentivirus overexpressing constitutively active human TGFβ2 (LV-TGFβ2). LV-TGFβ2-injected eyes, but not the contralateral eyes injected with a lentivirus containing a scrambled transgene (LV-Ctrl), exhibited significant IOP elevations one-week post-transduction (Figure 5*A*, Week 2, Δ_TGF-Ctrl_ = 4.0 mm Hg, *P* = 0.0143). By 2 weeks post-transfection, IOP in LV-TGFβ2 eyes reached 19.9 ± 4.7 mm Hg whereas IOP in LV-Control eyes remained at control levels (14.0 ± 1.2 mm Hg), with Δ_TGF-Ctrl_ = 5.9 mm Hg (*P* = 0.0002). IOP remained elevated throughout the 4 weeks after the injection (Week 5, Δ_TGF-Ctrl_ = 4.9 mm Hg, *P* = 0.0008). HC-06 (100 µM) microinjection into the anterior chamber of LV-TGFβ2 and LV-Ctrl eyes lowered IOP in LV-TGFβ2 eyes to 12.2 ± 1.7 mm Hg after 24 hours (Δ_postHC-preHC_ = -5.8 mm Hg) with no difference observed in IOP from LV-Ctrl eyes (12.6 ± 1.9 mm Hg, Δ_postHC-preHC_ = -0.3 mm Hg). LV-Ctrl eyes remained close to pre-injection levels post-HC-06 treatment (Figure 5*A*-*B*). IOP in LV-TGFβ2 eyes returned to hypertensive levels by 1-week post-HC-06 injection (Week 6-7, Δ_TGF-Ctrl_ = 3.9 mm Hg, *P* = 0.0201). To determine the effect of the bolus injection alone, LV-TGFβ2 and LV-Ctrl eyes were reinjected with PBS 2 weeks after re-establishing the OHT baseline. The sham injection transiently reduced IOP in LV-TGFβ2 (Δ_postPBS-prePBS_= -4.5 mm Hg) and LV-Ctrl (Δ_postPBSpre-PBS_= -1.2 mm Hg) eyes; however, LV-TGFβ2 eyes returned to hypertensive levels by 48 hours post-injection (Δ_TGF-Ctrl_ =3.6 mm Hg, p=0.0465) and to pre-injection levels after 72 hours (Δ_TGF-Ctrl_ =5.4 mm Hg, p=0.0002). Bolus injection was less effective than HC-06 at all time points 24 hours post-injection (Week 8-9, Figure 5*B*). These data indicate that selective pharmacological inhibition of TRPV4 effectively and reversibly blocks TGFβ2-induced OHT.

**Figure 5:**
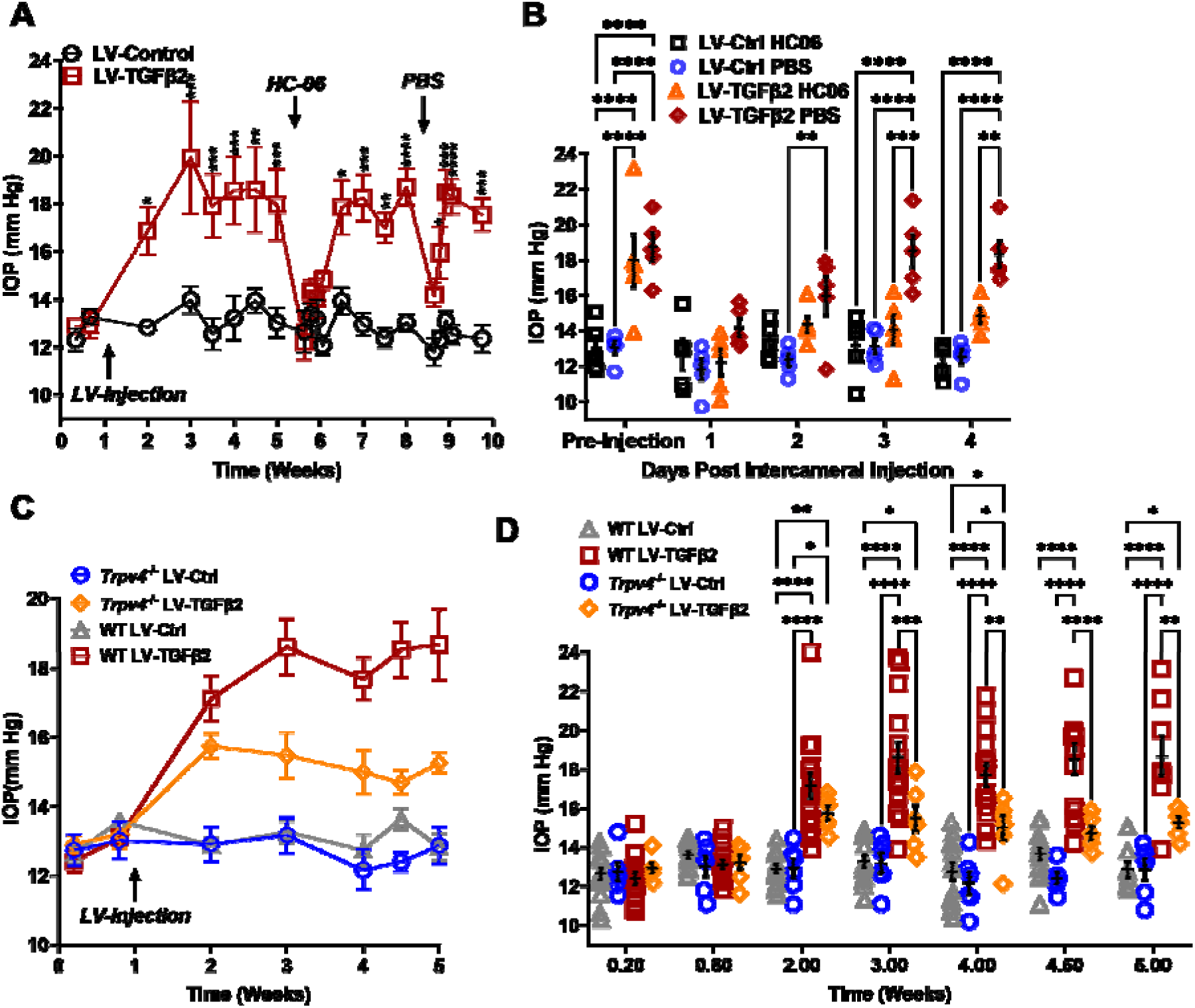
TRPV4 activation is necessary to maintain LV-TGFβ2-induced ocular hypertension. (**A**) Intravitreal injection of LV-TGFβ2 (week 1), but not LV-Control, elevates IOP in WT mice (N = 5 eyes/group) as early as one-week post-injection. Injection of TRPV4 antagonist HC-06, but not PBS, produced multiday IOP reduction in LV-TGFβ2 treated eyes. HC-06 and PBS injections did not affect IOP in LV-Control injected eyes. Two-way ANOVA with Bonferroni post-hoc analysis (**B**) Direct comparison of the results of PBS and HC-06 injections in the eyes shown in **A**. Two-way ANOVA with Bonferroni post-hoc analysis (**C**) Intravitreal injection of LV-TGFβ2 in *Trpv4^-/-^* mice (N = 6 eyes/group) resulted in only mild OHT; plotted against WT eyes at matching timepoints (3 WT cohorts including the 5 WT eyes shown in A-B, N = 8-15 eyes/group). (**D**) Statistical comparison of the IOP values shown in **C.** The IOP in LV-TGFβ2 WT eyes was significantly elevated compared to the LV-TGFβ2 *Trpv4^-/-^* eyes from 2 weeks post-injection. LV-Control injected eyes in WT or *Trpv4^-/-^* eyes remain close to the baseline value and are not significantly different. Two-way ANOVA with Bonferroni post-hoc analysis. (**A, C)** shows mean ± SEM. Data in (**B, D**) shows individual data points over mean ± SEM, *\* P* < 0.05, *\*\* P* < 0.01, *\*\*\* P* < 0.001, *\*\*\*\* P* < 0.0001

To further evaluate the TRPV4-dependence of TGFβ-induced OHT we took advantage of mice with global *Trpv4* knockdown (64–66). *Trpv4^-/-^* mice (N = 6) were injected with LV-TGFβ2 and LV-Ctrl vectors in contralateral eyes (Figure 5*C*). Additionally, two littermate control mice injected alongside the *Trpv4^-/-^*animals were added to previously collected WT LV-injected cohorts measured at the same timepoints (N = 8-15, Figure 5*C*). Pre-LV injection, IOP levels in *Trpv4*^-/-^ animals were comparable to the WT cohort, indicating that TRPV4 activity does not regulate normotension. Similarly, IOP in LV-Ctrl-injected eyes was not significantly different between WT and *Trpv4^-/-^* animals at any point in the experiment (peak Δ_CtrlKO-CtrlWT_ = -1.2 mm Hg, Figure 5*D*, SI Appendix Figure S5). By two weeks post-injection (Week 3), LV-TGFβ2-treated *Trpv4*^-/-^ eyes exhibited significantly lower IOP compared to the LV-TGFβ2 WT cohort (Δ_TGFKO-TGFWT_ = -3.1 mm Hg, *P* = 0.0009, Figure 5*C*). While LV-TGFβ2 injected *Trpv4^-/-^* eyes exhibited mild OHT, the effect was significantly reduced compared to WT eyes and IOP decreased by two weeks post-injection (Figure 5*C*-*D*).

### TGFβ2-induced and nocturnal OHT are non-additive but require TRPV4

Mammalian IOP is modulated by the circadian rhythm, with levels elevated at night and nocturnal IOP fluctuations implicated in POAG (7, 55, 67). We measured nocturnal (9:00-10:00 PM) IOP in LV-TGFβ2 (N = 4) and LV-Ctrl WT eyes (N = 4) from isoflurane anesthetized mice ∼ 2 months post-LV injection to determine whether nocturnal OHT is additive to TGFβ2-induced elevation observed during the daytime (12:00-2:00 PM, Figure 6*A*). LV-TGFβ2 injected eyes showed significant IOP elevation compared to LV-Ctrl eyes during daytime measurements (diurnal Δ_TGFβ-Ctrl_ = 7.9 mm Hg, *P* < 0.0001) but the difference vanished at night (nocturnal Δ_TGFβ-Ctrl_ = 0.2 mm Hg), indicating that TGFβ2-induced OHT is not additive to the circadian OHT. To test whether the IOP measurement was influenced by isoflurane-induced anesthesia, we repeated the nocturnal measurements in awake animals (N = 4-6 eyes/group). We observed no difference in nocturnal IOP between the two groups of animals was observed (Figure *6A-D*, SI Appendix Figure S6). To determine whether physiological (nocturnal) OHT requires TRPV4 we microinjected LV-Ctrl (N = 4) and hypertensive LV-TGFβ2 (N = 4) eyes with PBS or HC-06. PBS injection did not affect IOP in LV-Ctrl or LV-TGFβ2 eyes at day or night (Figure 6*E-F*) except for a single LV-TGFβ2 eye that exhibited abnormally high nocturnal IOP (37 mm Hg) at the four-day timepoint. Conversely, HC-06 injection blocked LV-TGFβ2-induced IOP during the day (*P* < 0.001) and significantly lowered IOP in LV-Ctrl and LV-TGFβ2 eyes at night (∼5 mm Hg; *P* < 0.01). These data indicate that i) TRPV4 activation is necessary for OHT in the TGFβ2 overexpression mouse model (Figures 5-6) and the circadian IOP elevations ii) TGFβ2 -evoked OHT does not affect nocturnal IOP elevation in mice, and iii) TRPV4 inhibition does not disrupt the mechanisms that maintain daytime normotensive IOP (Figures 5-6).

**Figure 6:**
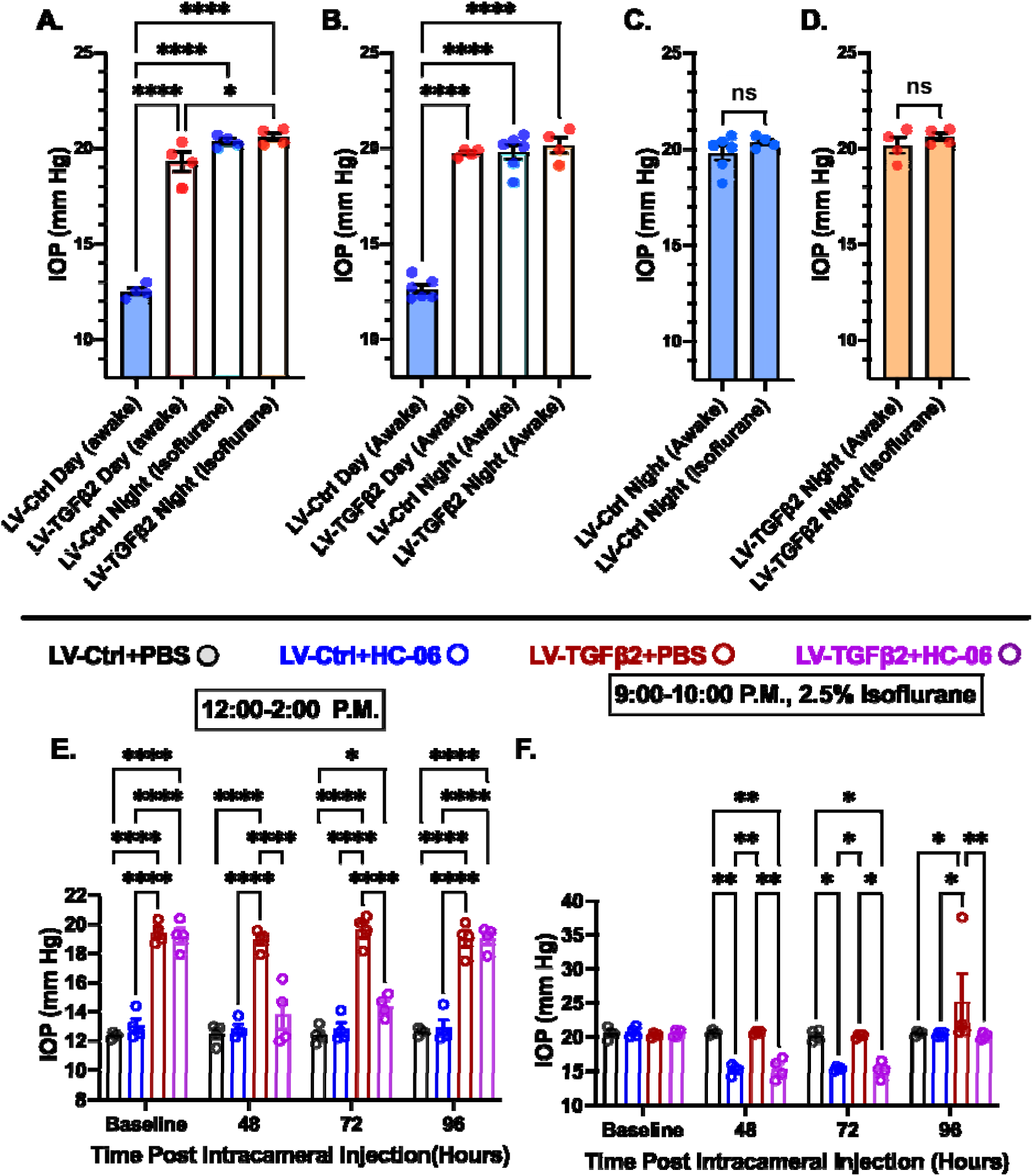
TRPV4 inhibition inhibits nocturnal IOP elevation in control and TGFβ2 overexpressing eyes. (**A-B**) Post-LV injection daytime (12-2:00 P.M) and nocturnal (9-10:00 P.M.) IOP compared in WT mice (N = 4-6 eyes/group) before drug treatment. LV-TGFβ2 eyes were elevated at daytime, but nocturnal OHT was not significantly different between LV-Ctrl and LV-TGFβ2 eyes in two separate cohorts of mice measured under isoflurane anesthesia (A) or while awake (B). (**C-D)** Anesthesia had no significant effect on measured nocturnal IOP. One-way ANOVA with Tukey’s multiple comparisons test. (**E**-**F)** PBS-injected eyes did not exhibit changes in daytime or nighttime intraocular pressure; however, HC-06 injection reduced TGFβ2-induced IOP elevations during the day and LV-Ctrl and LV-TGFβ2 nocturnal IOPs (N = 4 Eyes/Group); Two-way ANOVA with Bonferroni post-hoc analysis. Figures show datapoints over mean ± SEM, \**P* < 0.05*, ** P* < 0.01, *\*\*\* P* < 0.001, *\*\*\*\* P* < 0.0001

## Discussion

This study establishes a mechanistic framework that integrates biochemical and biomechanical risk factors of POAG and highlights the pivotal role of TRPV4, a polymodal Ca^2+^-permeable channel, as a key regulator of TM contractility and ocular hypertension. Our central finding is that the glaucoma-inducing cytokine TGFβ2 amplifies TRPV4 expression and activity, which in turn drives tonic increases in TRPV4 activation and TM contractility that are required to maintain elevated IOP. These observations position the pressure-activated channel as a molecular linchpin that links mechanical stress to neurodegeneration resulting from the obstruction of the primary outflow pathway. Our confirmation of the linkage between TGFβ signaling and TRPV4 in TM links POAG pathophysiology to fibrotic remodeling seen in other tissues (68, 69). Considering that current glaucoma treatments target secondary outflow mechanisms or incur side effects (such as hyperemia) (70, 71), the IOP lowering achieved through TRPV4 inhibition and gene knockdown promises a novel therapeutic avenue to mitigate ocular injury.

Glaucoma is a multifactorial disease with etiology that reflects the convergence of risk factors that include IOP and TGFβ2: epidemiological data correlate the incidence of POAG with the amplitude of IOP and [TGFβ2]_AH_ (22, 72), while chronic increases of either [TGFβ2]_i_ or IOP promote fibrotic remodeling of TM/SC and augment the flow resistance of the conventional pathway (17, 24, 25). TGFβ2-induced facility suppression has been historically attributed to changes in composition, crosslinking and amount of ECM (25, 26, 73, 74), activation of Hippo signaling and Rho kinase- (Rho/ROCK) mediated contractility (19, 57) and altered expression of genes encoding mitogen-activated protein kinase (MAPK), immune response, oxidative stress, and/or ECM pathways (75–77). Our discovery reveals bidirectional interplay between biomechanical and biochemical mechanisms: TGFβ2 impacts the expression and function of TM mechanosensors and *vice versa*, TRPV4 is required for TGFβ2-induced contractility: TGFβ2 (i) induced time-dependent upregulation of TRPV4 mRNA and amplified TRPV4-mediated calcium signaling, while (ii) TRPV4 was required to mediate TGFβ2 -induced TM hypercontractility and maintain chronic OHT in TGFβ2-treated mouse eyes. Microinjection of the selective antagonist HC-06 accordingly reduced IOP in LV- TGFβ2-treated eyes to baseline with hypotension persisting for ∼4 days and reversing to pre-injection OHT by day 7. The TRPV4-dependence of TGFβ2-induced OHT and contractility was corroborated in *Trpv4*^-/-^ mice and *in vitro* using TGFβ2-treated 3D hydrogel constructs. *In vivo*, pharmacological inhibition (∼100% reduction in OHT, transient) outperformed gene knockdown (∼50% reduction in OHT, stable), potentially due to compensatory mechanosensory mechanisms in Trpv4-*null* animals (55).

We’ve previously shown that TM TRPV4 is activated by physiological (5 – 25 mm Hg) pressure steps (39, 60) and (1 – 12%) strains (37, 41), which trigger downstream outflow-relevant signaling through Rho kinase, F-actin, tyrosine phosphorylation of FAK, paxillin and vinculin, lipid remodeling, and ECM release mechanisms (37, 41, 52). The present study extends those observations by revealing bidirectional effects of TRPV4 modulation on TM contractility (the agonist GSK101 induced, and the antagonist HC-06 suppressed, contractility in a 3D biomimetic model) and by establishing TRPV4 as an obligatory effector of OHT under physiological (circadian rhythmicity) and pathological conditions. These findings resolve conflicting reports of hypotensive vs. hypertensive effects of TRPV4 modulation (7, 36, 37, 39, 41, 52, 53, 55). Although TRPV4 activity has been suggested to lower IOP via phosphoinositide signaling in primary cilia (36), TM-resident endothelial nitric oxide synthase (eNOS) (7), release of polyunsaturated fatty acids (PUFAs) (53) and/or signaling downstream from Piezo1 mechanosensing (54), these mechanisms are challenged by evidence that TRPV4-regulated Ca^2+^ influx persists in TM cells with ablated primary cilia (37), eNOS expression in TM cells is vanishingly low (78–80), PUFAs such as arachidonic acid and EETs stimulate rather than inhibit TRPV4 (37) and TRPV4 signaling in TM cells is unaffected by Piezo1 inhibition and knockdown (39). Instead, the suppression of outflow facility by Piezo1 inhibitors applied under *in vitro* and *in vivo* conditions (39, 81) suggests that Piezo1 may oppose the hypertensive functions of TRPV4.

The TRPV4-dependence of increased outflow resistance is indicated by multiple lines of evidence. In biomimetic human TM scaffolds that support flow devoid of ciliary body or SC influences, TRPV4 inhibition enhanced, and activation suppressed, the facility (37). We accordingly demonstrate that the agonist (GSK101) induces contractility while the antagonist (HC-06) mitigates TGFβ2-induced hypercontractility and lowers IOP in TGFβ2-overexpressing eyes. Taken together, these results support a model wherein TRPV4-mediated pressure transduction is augmented by TGFβ2 to drive hypercontractility and fibrosis via Ca^2+^-and Rho-dependent stress fiber formation and reinforcement of focal ECM contacts (41, 82, 83) (Figure 7). TRPV4 is a thermosensitive ion channel with Q_10_ of ∼10 and optimal activation at physiological temperature of ∼34 - 37°C (84, 85). TGFβ2 -induced contractility of TM-populated hydrogels thus reflects both the severalfold transcriptional and functional amplification of TRPV4-mediated signaling (Figures 1-3) and constitutive temperature-facilitated activation of TRPV4 channels that is unmasked by the absence of hypercontractility in samples treated with HC-06. The modest contractility observed in HC-06-treated cells may reflect residual contributions from Piezo1, TRPC, and/or TREK-1 channels (37, 60, 86, 87). Parallels from studies conducted on heart, lung, liver, skin and articular cartilage similarly found that TRPV4 contributes to the progression of fibrosis induced by the cognate TGFβ1 (68, 88–91) and implicated the channel in bladder (92), heart (93, 94), and blood vessel (95) contractility. Indeed, conditional *Trpv4* ablation from smooth muscle cells lowered blood pressure (96, 97), an effect not dissimilar from IOP lowering in *Mgp:Trpv4* cKO mice (Figure 5).

**Figure 7:**
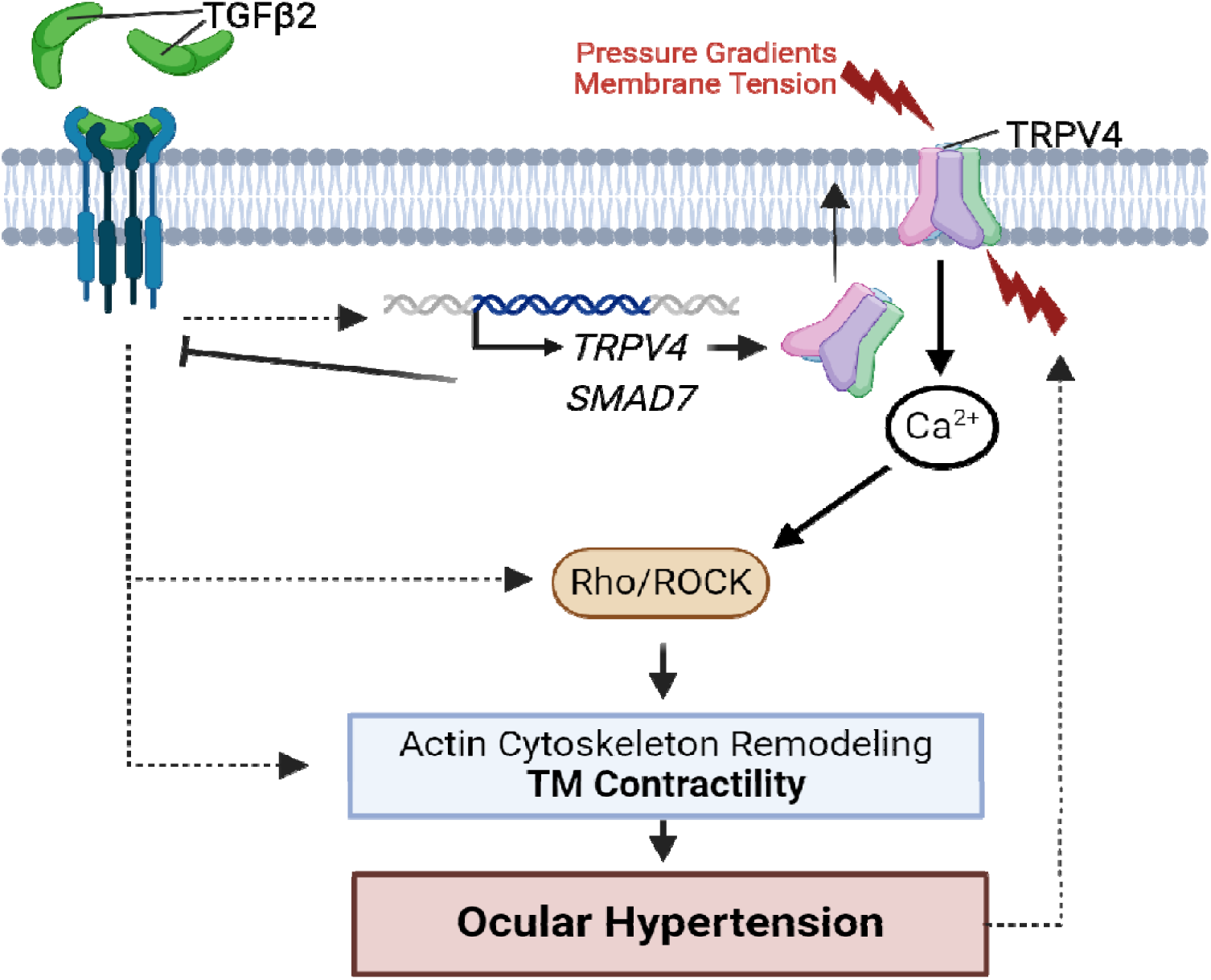
TGFβ2-TRPV4 interactions in TM remodeling and ocular hypertension. Chronic exposure to TGFβ2 induces upregulation of functional TRPV4 channels alongside the autoinhibitory canonical modulator SMAD7. TRPV4-mediated Ca^2+^ influx, canonical, and non-canonical TGFβ2 signaling stimulate the Rho/ROCK pathway to augment cytoskeletal contractility, and stimulate ECM release. Actomyosin contractility promotes outflow resistance and drives OHT and underpins a vicious feedforward TRPV4-dependent loop that maintains ocular hypertension.

TGFβ2-induced upregulation of *FN1, SNAIL1,* and *CTGF* transcripts (Figure 1) accords with RNA profiling studies which catalogued the cytokine’s role in transdifferentiation of TM cells into contractile myofibroblasts (25, 75, 98–103) whereas the decreased expression of *TGFΒR2* and increased abundance of SMAD7 mRNA indicate concurrent activation of autoinhibitory mechanisms associated with canonical TGFβ-family signaling (104). The-3-fold increase in TRPV4 transcription and responsiveness to GSK101, observed at POAG-relevant TGFβ2 concentrations (In AH ∼0.2-3.2 ng/ml; 20) showed a time course that mirrored facility reduction in human eyes treated with exogenous cytokine (105). A single 5 ng/ml TGFβ2 dose was sufficient to double the amplitude of the GSK101-evoked current and alter its rectification (Figure 3), with imaging experiments confirming robust and reproducible increases in Ca^2+^ signals across the 5/5 studied strains after 5 days of treatment (Figure 2). The effects of TGFβ2 on I_TRPV4_, membrane protein levels and [Ca^2+^]_GSK_ accord with increased expression of the TRPV4 gene. Precedents from other cell types (e.g., fibroblasts) suggest functional upregulation might involve increased trafficking of TRPV4- PI3Kg complexes and/or β-arrestin 1-dependent ubiquitination (106, 107). The upregulation of TRPV4/Piezo1 transcription by TGFβ2 predicts exaggerated responsiveness to mechanical loading, as reported for chemotherapy (108), neuropathic pain (109, 110), cancer (111), and diabetic neuropathy (112).

An important corollary of these findings is that they extend TRPV4’s role beyond pathology into the IOP homeostasis in healthy animals. TRPV4 inhibitors, ROCK and TM-specific expression of dominant negative scAAV2.*dnRhoA* constructs lower IOP across diverse OHT models (occlusion of the iridocorneal angle, TGFβ, glucocorticoids and the nocturnal cycle; 55, 62, 113). Physiological (circadian-induced) and pathological OHT were not additive (Figure 6) yet TRPV4 inhibition reversibly lowered nocturnal IOP in both untreated and TGFβ2 eyes to imply convergence at the level of TRPV4-Rho signaling as the final effector of OHT maintenance. Future studies should elucidate the mechanisms that mediate the reversibility of circadian TRPV4 activation. We conjecture potential involvement of the suprachiasmatic nucleus, the hypothalamus–pituitary–adrenal axis (114, 115), nocturnal release of norepinephrine and melatonin (116, 117) and altered TRPV4 modulation by β1 integrins (118), caveolin-1 (52), and cytoskeletal proteins (actin, actin adaptor proteins, microtubules) (119).

This study bridges biomechanical and biochemical paradigms of glaucomatous remodeling by extending the role of TGFβ2 beyond fibrosis to include TRPV4-dependent signaling and actomyosin contraction. We propose that TGFβ2 -induced upregulation of TRPV4 expression shifts the normotensive setpoint maintained by steady-state TRPV4, Piezo1 and TREK-1 activity (39, 60, 81) to heighten the TM sensitivity to mechanical cues, hijack contractile machinery and induce fibrosis that facilitates the pull of stress fibers on the increasingly “stiff” ECM (19). The absence of structural and functional visual phenotypes in TRPV4 KO mice (55, 66, 120) predicts that IOP lowering, suppression of fibrosis and protection of retinal neurons from pressure by small-molecule TRPV4 antagonists can take place without compromising homeostatic IOP regulation (70). The conserved TM TRPV4 expression (37) and TM physiology in mice vs. humans (121, 122) suggest that the findings reported might be relevant in a clinical context.

## Methods

### Animals

C57BL/6J mice were from JAX laboratories (Bar Harbor, ME), *Trpv4*^-/-^ (*Trpv4^tm1.1Ldtk^ Tg(KRT14-cre/ERT)20Efu/0;* MGI:5544606) mice were a gift from Wolfgang Liedtke (Duke University) (64, 65). The animals were maintained in a pathogen-free facility with a 12-hour light/dark cycle and ad libitum access to food and water, at a temperature of ∼22-23°C. Mice were 2-6 months in age prior to LV-injection; data from both male and female sexed animals were included in this study.

### Human TM Culture

De-identified postmortem eyes from donors with no history of glaucoma (pTM cells) were procured from Utah Lions Eye Bank with written informed consent of the donor’s families. TM cells were isolated from juxtacanalicular and corneoscleral regions as previously described (37, 39), in accordance with consensus characterization recommendations (123). pTM cells were cultured in Trabecular Meshwork Cell Medium (TMCM; Sciencell) in Collagen-I (Corning) coated culture flasks and glass coverslips at 37°C in a humidified atmosphere with 5% CO2. Fresh media was supplied every 2-3 days. Serum free (SF) media was mixed as needed by excluding fetal bovine serum (FBS, Sciencell) from the TMCM. A list of all pTM strains used is available in Table 1; all cells were used between passages 2-4. Cell lines were chosen based on availability at the time of experiments.

For contractility experiments pTM cells were isolated from healthy donor corneal rims discarded after transplant surgery, as previously described (19, 57, 124), and cultured according to established protocols (123, 125). Three pTM cell strains isolated from healthy donors and validated with dexamethasone-induced myocilin expression were used for contractility experiments. pTM cells were cultured in low-glucose Dulbecco’s Modified Eagle’s Medium (DMEM; Gibco; Thermo Fisher Scientific) containing 10% fetal bovine serum (FBS; Atlanta Biologicals) and 1% penicillin/streptomycin/glutamine (PSG; Gibco) and maintained at 37°C in a humidified atmosphere with 5% CO2. Fresh media was supplied every 2-3 days.

The experiments were conducted according to the tenets of the Declaration of Helsinki for the use of human tissue.

### Reagents

The TRPV4 antagonist HC-067047 (HC-06) was purchased from Millipore-Sigma or Cayman Biotech and dissolved in DMSO at 20mM. The TRPV4 agonist GSK1016790A (GSK101; Cayman Biotech) was dissolved in DMSO at 1mM. Aliquots were diluted into working concentrations (10-25 nM, GSK101; 5-100 µM, HC-06). Recombinant human TGFβ2 protein was ordered from R&D Systems and reconstituted in sterile 4 mM HCl with 0.1% BSA at 20 ug/mL.

### Quantitative Real-Time PCR

Gene-specific primers were used to detect the expression of target genes, as described (126). Total RNA was isolated using the Arcturus PicoPure RNA isolation kit (Thermofisher Scientific). cDNA was generated from total RNA using qScript XLT cDNA Supermix (Quanta Biosciences). SYBR Green based real-time PCR was performed with 2X GREEN Master Mix (Apex Bioresearch Products). Gapdh was used as an endogenous control to normalize fluorescence signals. Gene expression relative to GAPDH was measured using the comparative CT method (2^-[ΔCT(gene)- ΔCT(GAPDH)]^). All genes were assessed in 4-8 individual samples taken from 3-7 different pTM strains. The primer sequences, expected product length, and gene accession are given in Table 2.

**Table 2:**
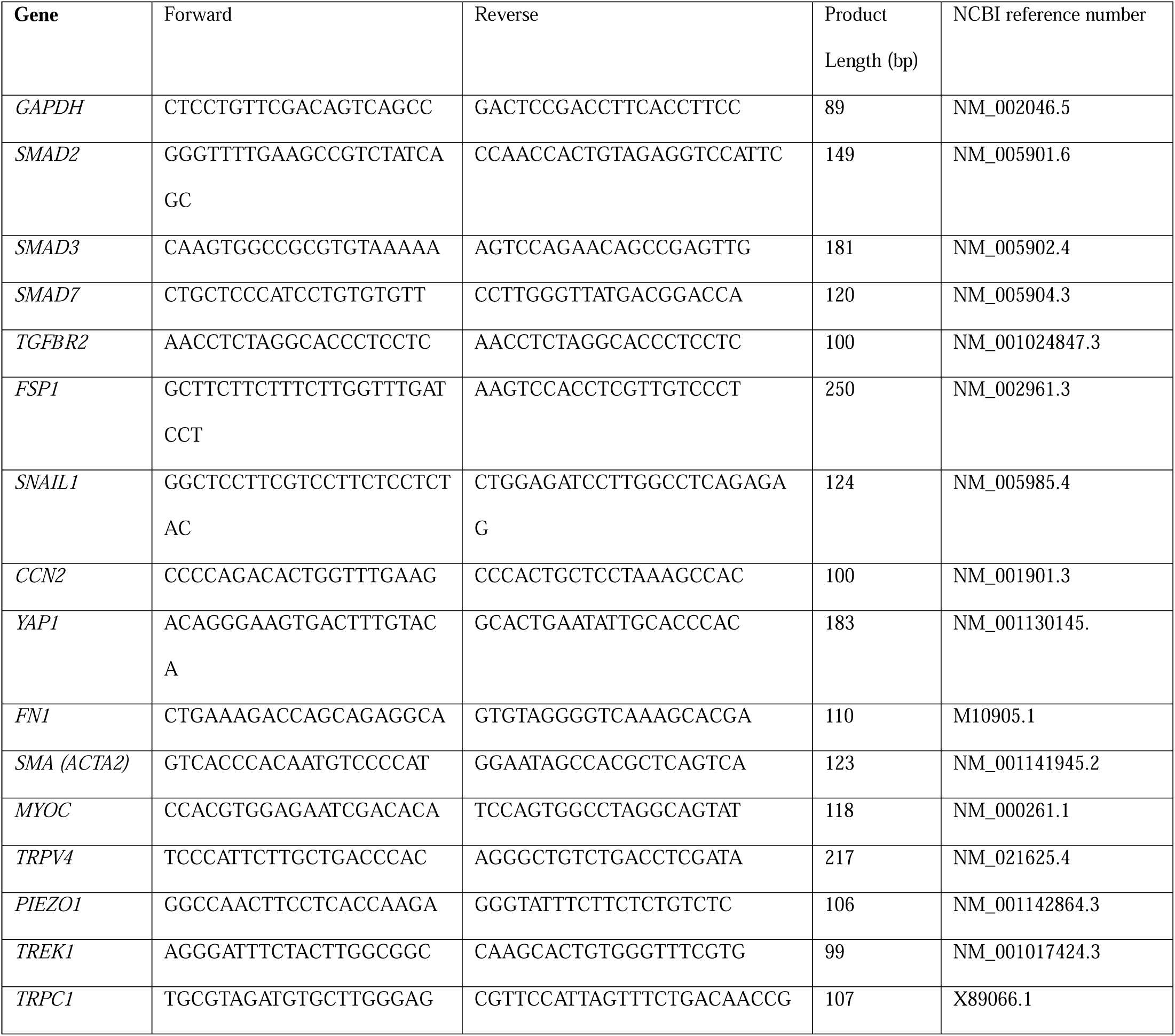
Sequences, product size, and reference numbers for PCR Primers used in this study.

### Western Blot

3 SF- or TGFβ2-treated samples were pelleted and pooled together from 3 different pTM samples within the same condition. To separate membrane proteins from heavier cellular debris the pooled cell pellets were homogenized in a hypotonic lysis buffer (20mM TRIS-HCl, 3mM MgCl2, 10mM NaCl, 10mM PMSF, 0.5 mM DTT, 20 mM NaF, 2 mM NaV, 0.5 µg/mL leupeptin) before centrifuging at 300x g for 5 minutes (4 °C). The resulting supernatant was removed and centrifuged again at >12,500 rpm for 30 minutes to pellet membrane proteins, which were then resuspended in RIPA Buffer (Santa Cruz). Proteins were separated on a 10% SDS-PAGE gel and transferred to polyvinylidene difluoride membranes (Bio-Rad). Membranes were blocked with 5% skim milk/2% BSA in TBST and incubated at 4 °C overnight with a primary antibody against TRPV4 (1:250, Alomone Labs #ACC-034) or rabbit antibody against β-tubulin (1:2000, Abcam #EPR1330). Appropriate secondary antibodies conjugated to HRP were used to visualize protein expression on an iBright CL750 imaging system (Thermo Fisher Scientific). β-Tubulin expression was used to standardize protein levels between samples.

### Calcium Imaging

pTM cells were seeded onto Collagen-I (Corning) coated coverslips and cultured in TMCM media (ScienCell) as described (39, 41). The cells were serum starved for 24 hours followed by serum-free TMCM with or without TGFβ2 (1 or 5 ng/mL) for 24 hours or five days. The cells were loaded with 10 µM of the ratiometric indicator Fura-2 AM (K_d_ at RT = 225 nM (Invitrogen/ThermoFisher) for 30-60 minutes. Coverslips were placed in a RC-26G chamber platform (Warner Instrument Corp) and perfused with external saline (pH 7.4) (in mM): 80 NaCl, 4.7 KCl, 1.2 MgCl2, 10 D-Glucose, 19.1 HEPES sodium salt, 2 CaCl_2_ and osmolality adjusted to 300 mOsm using D-mannitol. External solutions were delivered via a manually controlled gravity-fed eight-line manifold system, with perfusion speed kept constant to minimize changes in shear. Epifluorescence imaging was performed using an inverted Nikon Ti microscope with a 40x 1.3 N.A. oil objective and Nikon Elements AR software. 340 nm and 380 nm excitation were delivered by a high-intensity 150W Xenon arc lamp (Lambda DG-4; Sutter Instruments), high pass-filtered at 510 nm and detected with a 12-bit Delta Evolve camera (Photometrics/Teledyne). GSK101 (10 nM) evoked Δ[Ca^2+^]_i_ was assessed as ΔR/R (dividing the difference between the peak GSK-evoked F_340_/F_380_ signal during stimulation and baseline F_340_/F_380_ signal by the baseline F_340_/F_380_ signal). Every data point represents a separate experimental day and pTM cell strain, each with 3-5 control and 3-5 TGFβ2-treated slides tested on the same day. TGFβ2 datapoints represent the average GSK101 evoked ΔR/R across all TGFβ2 cells as a % of the average ΔR/R of control cells from the same cell strain on the same day.

### Collagen hydrogel contraction assay

Rat tail collagen type I (Corning, Thermo Fisher Scientific) was prepared at a concentration of 1.5 mg/ml according to the manufacturer’s instructions. Five hundred microliters of the hydrogel solution were pipetted into 24-well culture plates. Upon complete collagen polymerization, pTM cells were seeded at 1.5 × 105 cells/well atop the hydrogels and cultured in DMEM + 10% FBS + 1% PSG for 48 hours to facilitate even cell spreading. Next, constructs were cultured in serum-free DMEM + 1% PSG supplemented with: i) control (vehicle: 0.008 mM HCl + 0.0004% BSA; 0.025% DMSO), ii) TGFβ2 (5 ng/ml; R&D Systems), or iii and iv) TGFβ2 + HC067047 (5 µM in DMSO) for 36 hours before carefully releasing the hydrogels from the walls using a sterile 10 µl pipette tip to facilitate contraction. The next morning, fresh serum-free DMEM + 1% PSG supplemented with 0.0025% DMSO (= vehicle) was added to groups i-iii; group iv received serum-free DMEM + 1% PSG supplemented with GSK1016790A (25 nM in DMSO). Plates were longitudinally imaged at 600 dpi resolution with a CanoScan LiDE 300 flatbed scanner (Canon USA) at 0, 15, 30, 60, and 120 minutes. Hydrogel construct size was quantified using FIJI software (National Institutes of Health) (127).

### Electrophysiology

Borosilicate patch-clamp pipettes (WPI) were pulled using a P-2000 horizontal micropipette puller (Sutter Instruments), with a resistance of 6-8 MΩ. The internal solution contained (mM): 125 K-gluconate, 10 KCl, 1.5 MgCl2, 10 HEPES, 10 EGTA, pH 7.4. Patch clamp data was acquired with a Multiclamp 700B amplifier, pClamp 10.6 software and Digidata 1440A interface (Molecular Devices), sampled at 5kHz and analyzed with Clampfit 10.7. Current-voltage relationships were assessed using V_m_ steps from -100 to + 100 mV against a holding potential of -30 mV. Current density was measured as the average current during GSK101 exposure subtracted by the average current from the same cell during baseline perfusion.

### IOP Measurements

A TonoLab rebound tonometer (Colonial Medical Supply) was used to measure IOP of awake mice between 12-2 P.M. IOP was determined from the mean of 10-20 tonometer readings. Nocturnal measurements were conducted between 9-10 P.M. in awake animals or under 2.5% isoflurane delivered by a Somnosuite isoflurane vaporizer (Kent Scientific). After animals recovered from intracameral HC-06/PBS injections, IOP was measured daily. IOP was measured every day for 4-5 consecutive days to confirm a stable return to baseline. IOP data for individual cohorts was binned into weeks of experimental time to group values for analysis.

### Lentiviral Transduction

Lentiviral stock for TGFβ2 (C226,228S) was purchased from VectorBuilder Inc. (VB170816-1094fnw, pLV[Exp]-CMV> {hTGFB2[NM_003238.3](C226,228S)}) (23). Scrambled control lentivirus was purchased from SignaGen Laboratories (LM-CMV-Null-Puro). Mice were anesthetized with an intraperitoneal IP injection of ketamine/xylazine (90 mg/10 mg/ kg body weight), followed by eyedrops containing 0.5% proparacaine hydrochloride and 1% tropicamide ophthalmic solution to numb the eyes and dilate the pupils. Anesthetized mice were secured to allow stereotaxic injection of lentivirus. Intravitreal injections were conducted by creating a guide hole with a 30-gauge needle 1-2 mm equatorial of the cornea-scleral border followed by insertion of a 12° beveled 33-gauge Hamilton syringe (Hamilton Company) secured to a stereotaxic rig (World Precision Instruments) used to insert the needle 2-3 mm into the eye. Each eye was injected with a 2uL bolus of lentivirus diluted to 1×10^6^ TU/µL over the course of one minute, before the needle was quickly drawn and the pilot hole treated with erythromycin ophthalmic ointment USP (Bausch & Lomb). The efficiency of LV-TGFβ2 OHT induction in WT animals was close to 100%. No differences in observable health post-injection were detected between wild type and *Trpv4*^-/-^ animals or LV-Ctrl and LV-TGFβ2 injected animals.

### Intracameral Microinjections

Mice were anesthetized and treated with eyedrops as above, before being placed on an isothermal heating pad. HC-06 (100 µM) or PBS with DMSO (0.5%) as a vehicle were injected into the anterior chamber using a blunt tip Hamilton syringe (Hamilton Company) through a guide hole made using a 30-gauge needle. At the end of each injection a small air bubble was introduced to seal the cornea and minimize fluid outflow. 0.5% Erythromycin ophthalmic ointment USP (Bausch & Lomb) was applied to the eye after the procedure. Intracameral injections were not associated with observable inflammation, corneal opacity or behavioral changes. For the nocturnal IOP experiments in Figure 6, both eyes of two animals were injected with PBS while two were injected with HC-06. When OHT was stably reestablished a week post-injection, the treatment groups were switched, and experiments repeated resulting in four eyes/treatment group for Figure 6*E-F*.

### Statistical Analysis

GraphPad Prism 9 was used for statistical analysis. Means are plotted ± SEM unless otherwise noted. One-sample t-tests were used to determine whether TGFβ2 treated groups were significantly different than untreated control groups, while one-way ANOVA or two-way ANOVA along with Tukey or Bonferroni’s multiple comparisons test were used to compare multiple groups.

### Study Approval

The animal experimental protocols were conducted in accordance with the NIH Guide for the Care and Use of Laboratory Animals and the ARVO Statement for the Use of Animals in Ophthalmic and Vision Research and were approved by the Institutional Animal Care and Use Committee at the University of Utah.

## Data Availability

Individual datapoints for in-vivo figures, and unedited/uncropped annotated western blot images, are included in the supplementary data files for this manuscript. Further information about the data presented in this manuscript is available from the corresponding authors upon reasonable request.

## Author Contributions

C.N.R. and D.K. designed the primary research study. C.N.R., M.L., A.S., S.N.R., D.K., Y.T.T., M.L. performed research, C.N.R., S.N.R., Y.T.T., S.H., D.K. analyzed the data, and C.N.R. and D.K. wrote the paper.

## Acknowledgements & Funding

We thank Dr. Paloma Liton (Duke University) for the generous gift of LV-TGFβ2^C226,228S^ lentivirus stock during the pilot stages of this experiment, and Dr. Gulab Zode for the availability of the LV-TGFβ2^C226,228S^ construct on Vectorbuilder. We additionally thank Dr. Wolfgang Liedtke (Duke University and Regeneron) for *Trpv4^−/−^* mice.

The study was supported by the National Institutes of Health (T32EY024234 to CNR and DK, R01EY022076, R0EY1031817, P30EY014800 to DK, R01EY034096 to SH, R01EY022359, R01EY005722 to WDS), Crandall Glaucoma Initiative, Stauss-Rankin Foundation, and Unrestricted Grants from Research to Prevent Blindness to Ophthalmology Departments at the University of Utah and Duke University.

Schematics were made using Biorender.com.

## Supplemental Information

**Supplementary Figure S1:**
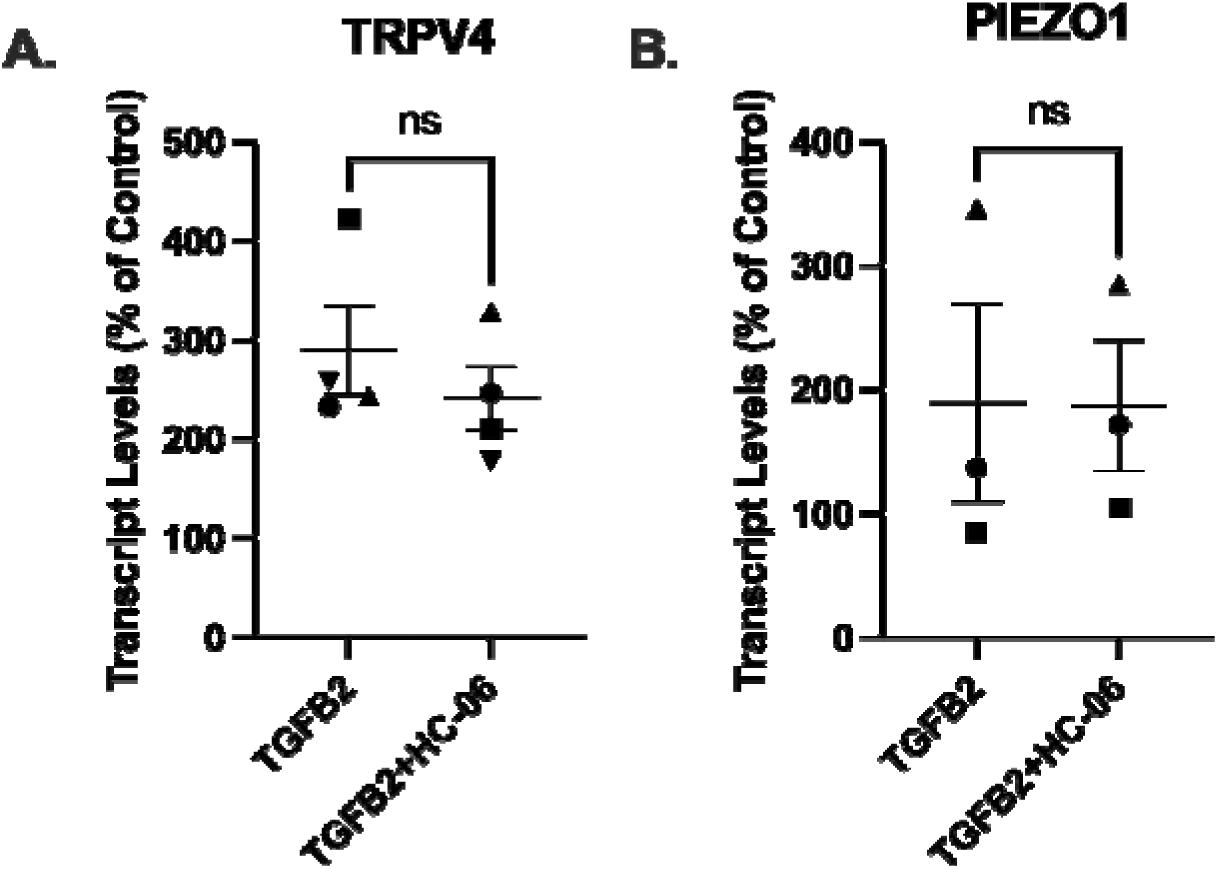
No significant difference was seen in *TRPV4* or *PIEZO1* expression between pTM samples treated with TGFβ2 (1 ng/mL) alone or TGFβ2 + TRPV4 antagonist HC-06 (5µM) for five days. N=3-4 independent experiments. Within each gene, symbols indicate paired samples. Wilcoxon matched-pairs signed rank test and paired t-test used respectively.

**Supplementary Figure S2:**
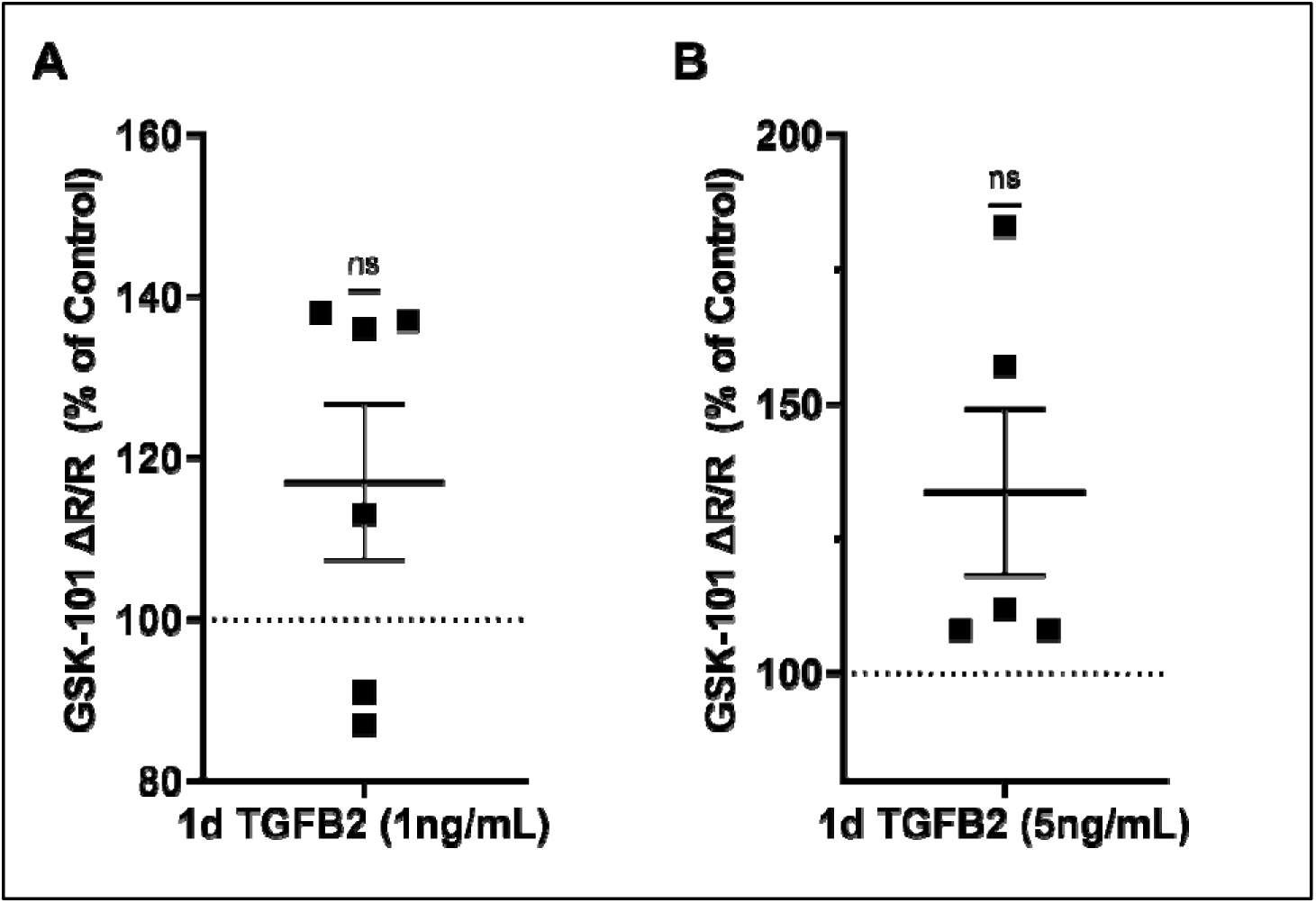
TGFβ2 concentrations of 1 ng/mL (**A)** and 5 ng/mL (**B)** did not significantly increase TRPV4-induced calcium influx with respect to control cells. Individual statistical analysis of experiments shown in Fig. 3*A* (1 ng/mL: *P* = 0.138, 5ng/mL: *P* = 0.095), one sample t-test.

**Supplementary Figure S3:**
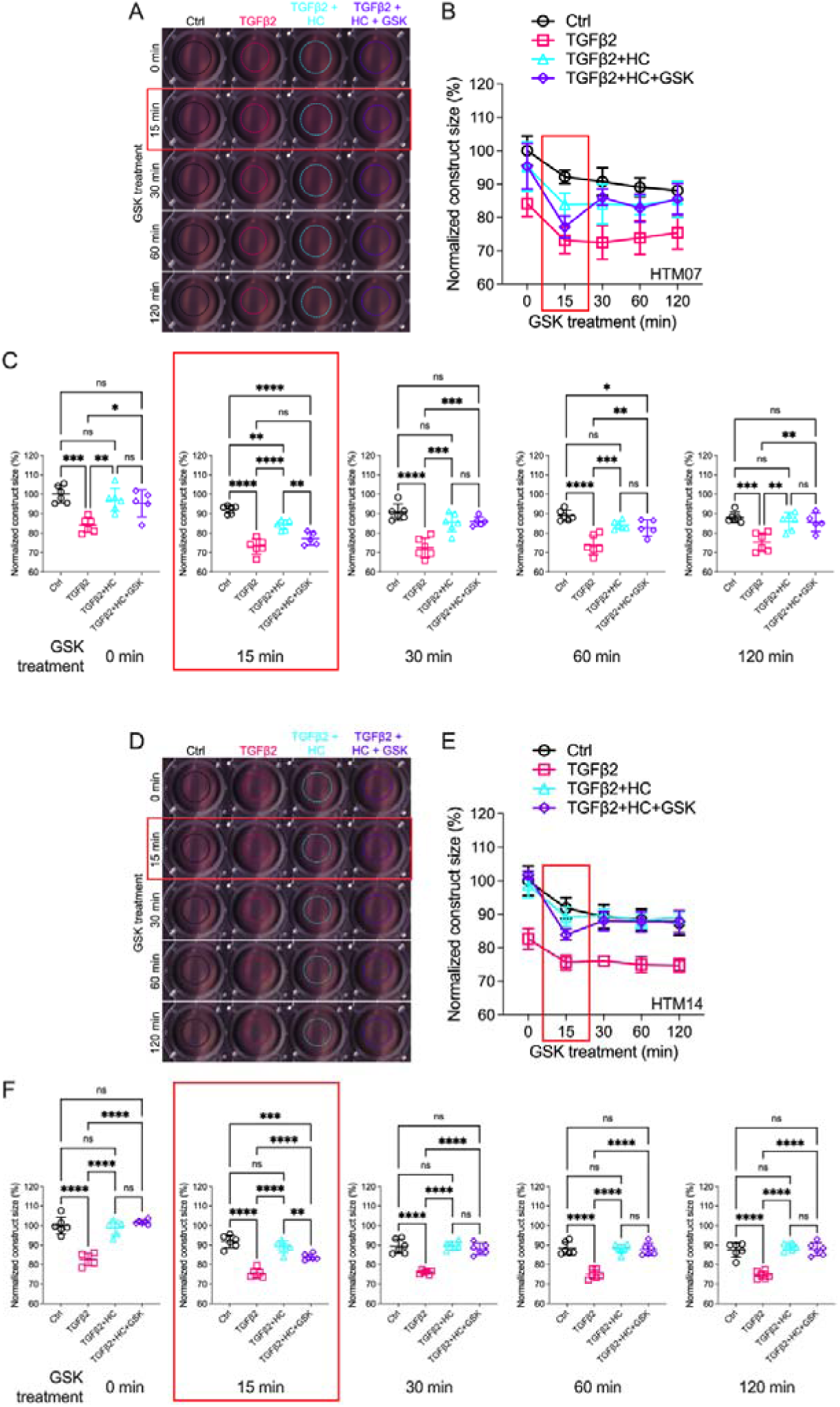
TRPV4 activation is obligatory for TGFβ2-induced TM cell contractions. (**A, D**) Representative longitudinal 24-well plate scans of collagen type I hydrogels seeded with two distinct pTM strains (pTM 1: **A**-**C**, pTM 2: **D**-**F**) subjected to the different treatments as in Fig. 4. (**B, E**) Longitudinal quantification of hydrogel construct size. (**C, F**) Detailed comparisons between groups at each experimental time point (N = 6 experimental replicates/ pTM strain). One-way ANOVA with Tukey multiple comparisons test, data in (**B, D**) shows individual data points over mean ± SEM *\* P* < 0.05, *\*\* P* < 0.01, *\*\*\* P* < 0.001, *\*\*\*\* P* < 0.0001.

**Supplementary Figure S4:**
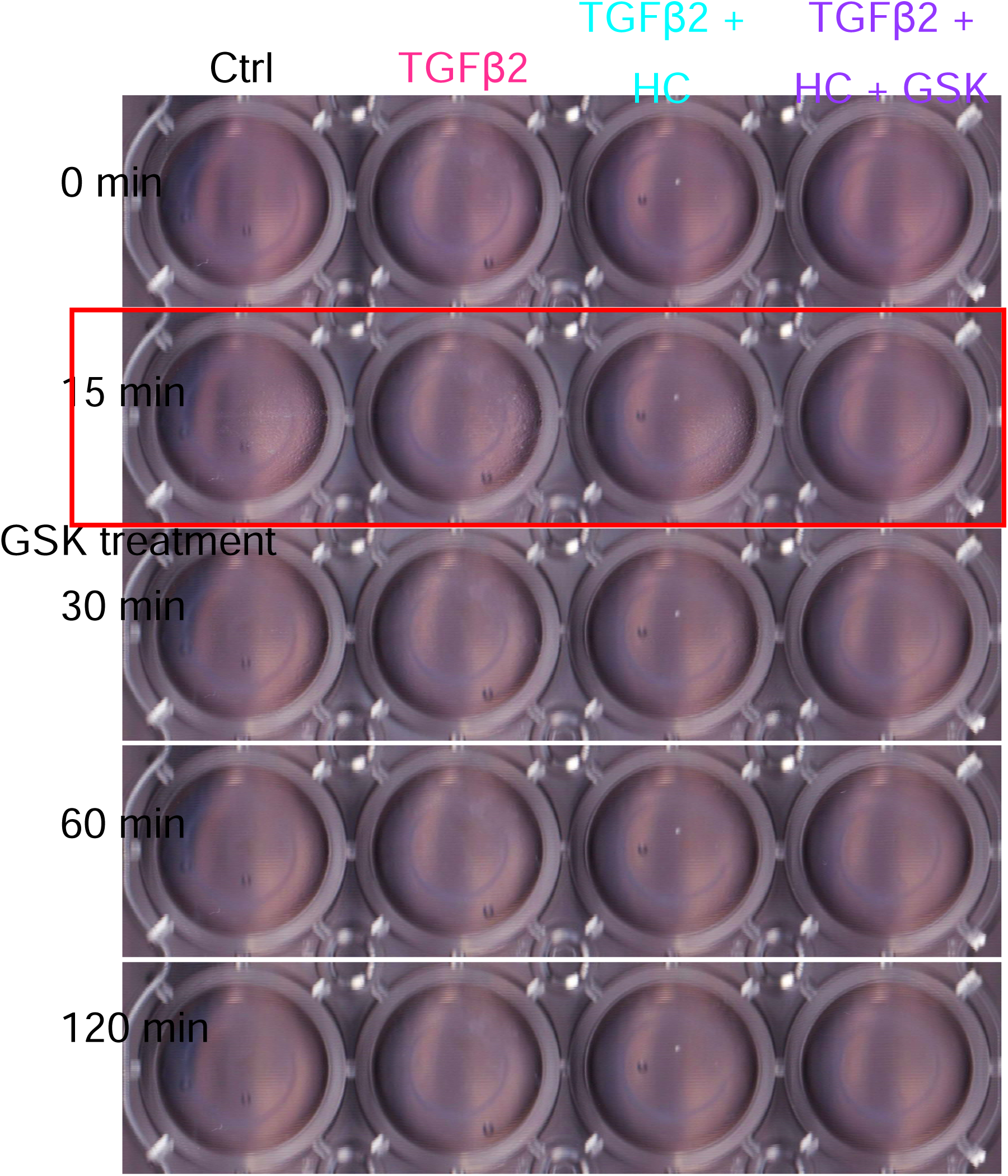
Detailed view of pTM seeded collagen constructs used in Figure 4 and S3. High resolution representative image of collagen gels used for contractility experiments without circle around periphery of gel.

**Supplementary Figure S5:**
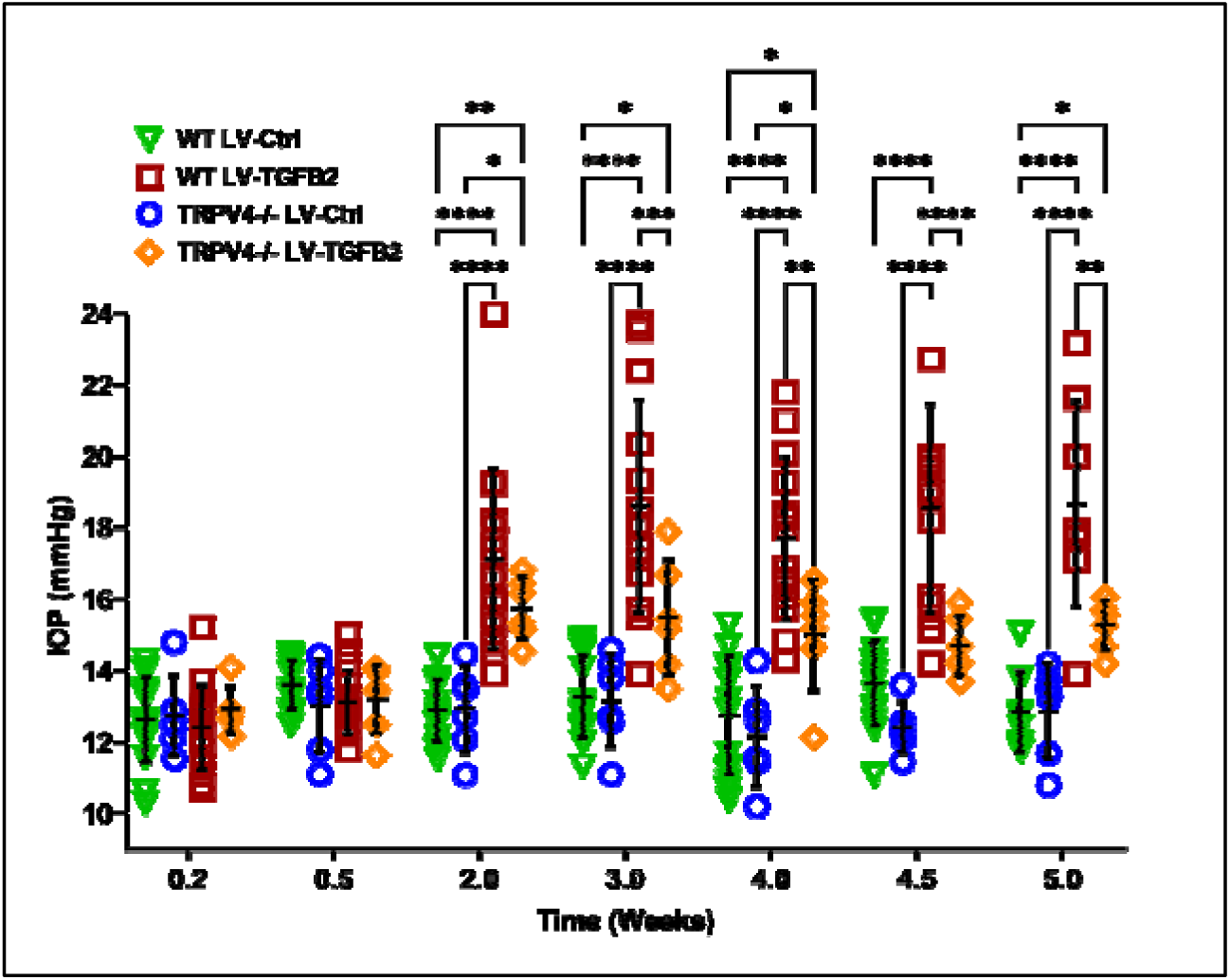
Expansion of Fig. 5*D*. IOP in LV-TGFβ2-injected eyes was significantly elevated compared to both LV-Ctrl injected WT and *Trpv4*^-/-^ eyes, as well as LV-TGFβ2-injected *Trpv4*^-/-^ eyes (N = 6 eyes/condition).

**Supplementary Figure S6:**
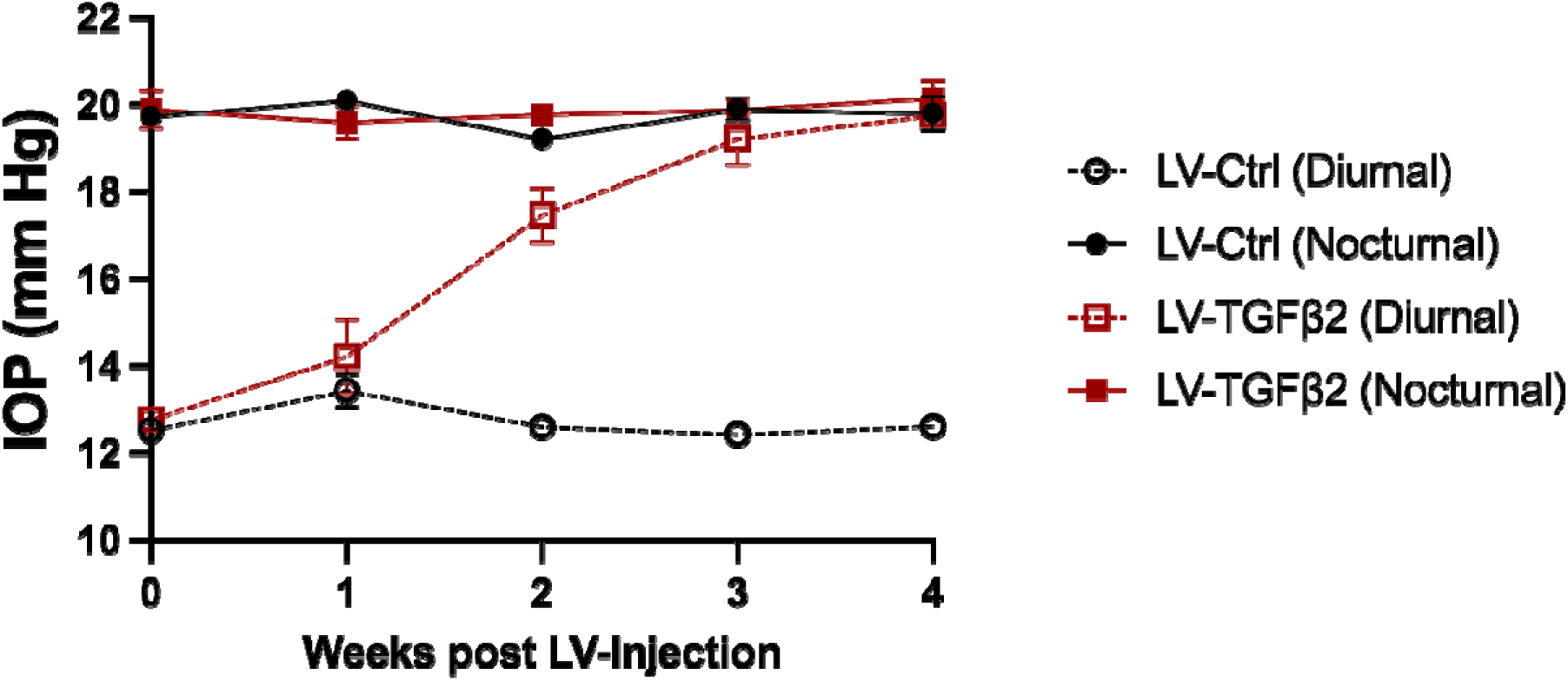
Nocturnal IOP is not significantly affected by LV-TGFβ2 overexpression. Expanded time series of IOP measured weekly from a second cohort of mouse eyes (Fig. 6B) injected with LV-Ctrl (n = 6 eyes) or LV-TGFβ2 (n = 4 eyes). LV-TGFβ2 resulted in elevated diurnal IOP which gradually approached the IOP seen in nocturnal measurements, but did not further elevate IOP above nocturnal values. In this cohort, both diurnal and nocturnal measurements were made in awake animals.

**Supplementary Information.**
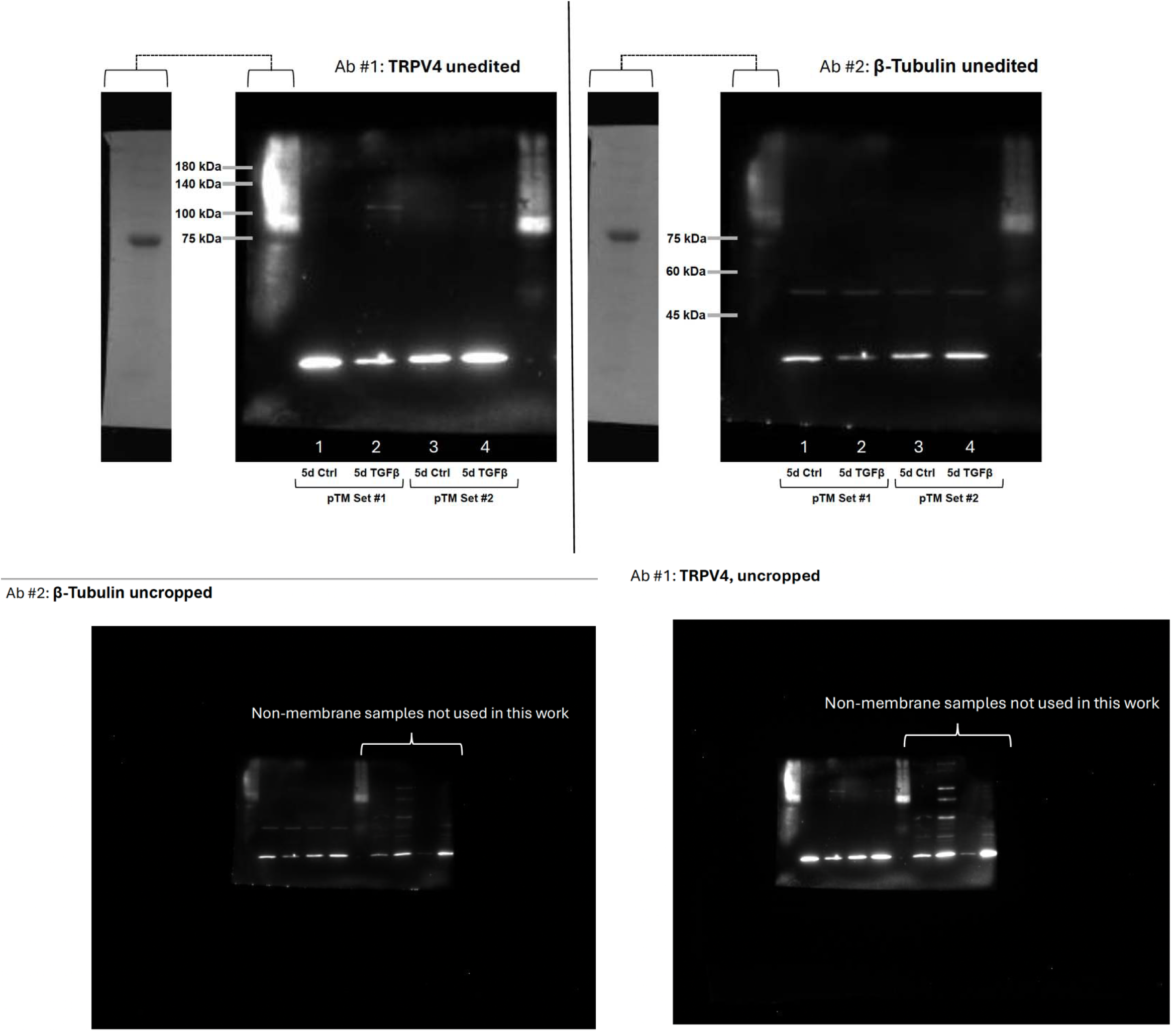
Uncropped Western blots.

## Notes

***COI***: The authors declare no conflict of interest, financial or otherwise. D.K. is a cofounder of TMClear and co-inventor of patents (US 2015/0133411, US20230026696) related to the development of cornea-permeant TRPV4 channel antagonists. The patents were licensed to TMClear by the University of Utah.

### Competing Interest Statement

The authors have declared no competing interest.

### Summary of Updates

OP measurements was conducted in another cohort of TGFB2-treated mice under diurnal and nocturnal conditions.

## References

1. Y. C. Tham, et al., Global prevalence of glaucoma and projections of glaucoma burden through 2040: A systematic review and meta-analysis. Ophthalmology 121, 2081–2090 (2014).

2. M. O. Gordon, et al., The Ocular Hypertension Treatment Study: Baseline factors that predict the onset of primary open-angle glaucoma. Arch. Ophthalmol. 120, 714–720 (2002).

3. A. Heijl, et al., Reduction of intraocular pressure and glaucoma progression: Results from the Early Manifest Glaucoma Trial. Arch. Ophthalmol. 120, 1268–1279 (2002).

4. M. Almasieh, A. M. Wilson, B. Morquette, J. L. Cueva Vargas, A. Di Polo, The molecular basis of retinal ganglion cell death in glaucoma. Prog. Retin. Eye Res. 31, 152–181 (2012).

5. C. Baudouin, M. Kolko, S. Melik-Parsadaniantz, E. M. Messmer, Inflammation in Glaucoma: From the back to the front of the eye, and beyond. Prog. Retin. Eye Res. 83 (2021).

6. B. A. Ellingsen, W. M. Grant, The relationship of pressure and aqueous outflow in enucleated human eyes. Invest. Ophthalmol. 10, 430–437 (1971).

7. P. D. Patel, et al., Impaired TRPV4-eNOS signaling in trabecular meshwork elevates intraocular pressure in glaucoma. Proc. Natl. Acad. Sci. U. S. A. 118 (2021).

8. A. W. De Kater, A. Shahsafaei, D. L. Epstein, Localization of smooth muscle and nonmuscle actin isoforms in the human aqueous outflow pathway. Investig. Ophthalmol. Vis. Sci. 33, 424–429 (1992).

9. David Krizaj, “No cell is an island: trabecular meshwork ion channels as sensors of the ambient milieu” in Glaucoma Research and Clinical Advances: 2020 to 2022, Vol. 3, J. R. Samples, P. A. Knepper, Eds. (Kugler Publications, 2020), pp. 1–10.

10. J. M. Baumann, et al., TRPV4 and chloride channels mediate volume sensing in trabecular meshwork cells. Am. J. Physiol. Physiol. (2024). 10.1152/ajpcell.00295.2024.

11. R. F. Brubaker, The effect of intraocular pressure on conventional outflow resistance in the enucleated human eye. Invest. Ophthalmol. 14, 286–292 (1975).

12. A. Karimi, et al., The Effect of Intraocular Pressure Load Boundary on the Biomechanics of the Human Conventional Aqueous Outflow Pathway. Bioengineering 9 (2022).

13. T. Borrás, Gene expression in the trabecular meshwork and the influence of intraocular pressure. Prog. Retin. Eye Res. 22, 435–463 (2003).

14. R. F. Ramos, G. M. Sumida, W. Daniel Stamer, Cyclic mechanical stress and trabecular meshwork cell contractility. Investig. Ophthalmol. Vis. Sci. 50, 3826–3832 (2009).

15. C. Luna, G. Li, P. B. Liton, D. L. Epstein, P. Gonzalez, Alterations in gene expression induced by cyclic mechanical stress in trabecular meshwork cells. Mol. Vis. 15, 534–544 (2009).

16. T. S. Acott, J. A. Vranka, K. E. Keller, V. K. Raghunathan, M. J. Kelley, Normal and glaucomatous outflow regulation. Prog. Retin. Eye Res. 82 (2021).

17. R. Fuchshofer, E. R. Tamm, The role of TGF-β in the pathogenesis of primary open-angle glaucoma. Cell Tissue Res. 347, 279–290 (2012).

18. M. A. Johnstone, Intraocular pressure regulation: Findings of pulse-dependent trabecular meshwork motion lead to unifying concepts of intraocular pressure homeostasis. J. Ocul. Pharmacol. Ther. 30, 88–93 (2014).

19. H. Li, V. K. Raghunathan, W. D. Stamer, P. S. Ganapathy, S. Herberg, Extracellular Matrix Stiffness and TGFβ2 Regulate YAP/TAZ Activity in Human Trabecular Meshwork Cells. Front. Cell Dev. Biol. 10 (2022).

20. P. Agarwal, A. M. Daher, R. Agarwal, Aqueous humor TGF-β2 levels in patients with open-angle glaucoma: A meta-analysis. Mol. Vis. 21, 612–620 (2015).

21. Y. Ochiai, H. Ochiai, Higher concentration of transforming growth factor-β in aqueous humor of glaucomatous eyes and diabetic eyes. Jpn. J. Ophthalmol. 46, 249–253 (2002).

22. R. C. Tripathi, J. Li, W. F. A. Chan, B. J. Tripathi, Aqueous humor in glaucomatous eyes contains an increased level of TGF-β2. Exp. Eye Res. 59, 723–728 (1994).

23. S. V. Patil, R. B. Kasetti, J. C. Millar, G. S. Zode, A Novel Mouse Model of TGFβ2-Induced Ocular Hypertension Using Lentiviral Gene Delivery. Int. J. Mol. Sci. 23 (2022).

24. A. R. Shepard, et al., Adenoviral gene transfer of active human transforming growth factor-β2 elevates intraocular pressure and reduces outflow facility in rodent eyes. Investig. Ophthalmol. Vis. Sci. 51, 2067–2076 (2010).

25. D. L. Fleenor, et al., TGFβ2-induced changes in human trabecular meshwork: Implications for intraocular pressure. Investig. Ophthalmol. Vis. Sci. 47, 226–234 (2006).

26. M. Montecchi-Palmer, et al., TGFβ2 induces the formation of cross-linked actin networks (CLANs) in human trabecular meshwork cells through the smad and non-smad dependent pathways. Investig. Ophthalmol. Vis. Sci. 58, 1288–1295 (2017).

27. R. K. Coker, et al., Transforming growth factors-β1, -β2, and -β3 stimulate fibroblast procollagen production in vitro but are differentially expressed during bleomycin-induced lung fibrosis. Am. J. Pathol. 150, 981–991 (1997).

28. B. Santiago, et al., Topical application of a peptide inhibitor of transforming growth factor-β1 ameliorates bleomycin-induced skin fibrosis. J. Invest. Dermatol. 125, 450–455 (2005).

29. Y. Yue, K. Meng, Y. Pu, X. Zhang, Transforming growth factor beta (TGF-β) mediates cardiac fibrosis and induces diabetic cardiomyopathy. Diabetes Res. Clin. Pract. 133, 124–130 (2017).

30. Y. Zhang, et al., Overexpression of TGF-b1 induces renal fibrosis and accelerates the decline in kidney function in polycystic kidney disease. Am. J. Physiol. - Ren. Physiol. 319, F1135–F1148 (2020).

31. M. Walker, M. Godin, A. E. Pelling, Mechanical stretch sustains myofibroblast phenotype and function in microtissues through latent TGF-β1 activation. Integr. Biol. 12, 199–210 (2020).

32. P. J. Wipff, D. B. Rifkin, J. J. Meister, B. Hinz, Myofibroblast contraction activates latent TGF-β1 from the extracellular matrix. J. Cell Biol. 179, 1311–1323 (2007).

33. G. Zhen, et al., Mechanical stress determines the configuration of TGFβ activation in articular cartilage. Nat. Commun. 12 (2021).

34. N. E. Cabrera-Benítez, et al., Mechanical stress induces lung fibrosis by epithelial-mesenchymal transition. Crit. Care Med. 40, 510–517 (2012).

35. S. C. Wei, J. Yang, Forcing through Tumor Metastasis: The Interplay between Tissue Rigidity and Epithelial-Mesenchymal Transition. Trends Cell Biol. 26, 111–120 (2016).

36. N. Luo, et al., Primary cilia signaling mediates intraocular pressure sensation. Proc. Natl. Acad. Sci. U. S. A. 111, 12871–12876 (2014).

37. D. A. Ryskamp, et al., TRPV4 regulates calcium homeostasis, cytoskeletal remodeling, conventional outflow and intraocular pressure in the mammalian eye. Sci. Rep. 6 (2016).

38. Y. F. Yang, Y. Y. Sun, D. M. Peters, K. E. Keller, The Effects of Mechanical Stretch on Integrins and Filopodial-Associated Proteins in Normal and Glaucomatous Trabecular Meshwork Cells. Front. Cell Dev. Biol. 10 (2022).

39. O. Yarishkin, et al., Piezo1 channels mediate trabecular meshwork mechanotransduction and promote aqueous fluid outflow. J. Physiol. 599, 571–592 (2021).

40. J. P. M. White, et al., TRPV4: Molecular conductor of a diverse orchestra. Physiol. Rev. 96, 911–973 (2016).

41. M. Lakk, D. Krizaj, TRPV4-Rho signaling drives cytoskeletal and focal adhesion remodeling in trabecular meshwork cells. Am. J. Physiol. - Cell Physiol. 320, C1013–C1030 (2021).

42. V. Katari, N. Bhavnani, S. Paruchuri, C. Thodeti, TRPV4 regulates matrix stiffness-dependent activation of YAP/VEGFR2 signaling via Rho/Rho kinase/LATS1/2 pathway. FASEB J. 35 (2021).

43. B. Nilius, T. Voets, The puzzle of TRPV4 channelopathies. EMBO Rep. 14, 152–163 (2013).

44. M. L. Thibodeau, et al., Compound heterozygous TRPV4 mutations in two siblings with a complex phenotype including severe intellectual disability and neuropathy. Am. J. Med. Genet. Part A 173, 3087–3092 (2017).

45. C. J. Klein, et al., TRPV4 mutations and cytotoxic hypercalcemia in axonal Charcot-Marie-Tooth neuropathies. Neurology 76, 887–894 (2011).

46. Z. Daneva, et al., Caveolar peroxynitrite formation impairs endothelial TRPV4 channels and elevates pulmonary arterial pressure in pulmonary hypertension. Proc. Natl. Acad. Sci. U. S. A. 118 (2021).

47. O. Pochynyuk, O. Zaika, R. G. O’Neil, M. Mamenko, Novel insights into TRPV4 function in the kidney. Pflugers Arch. Eur. J. Physiol. 465, 177–186 (2013).

48. M. W. G. Roberts, et al., TRPV4 receptor as a functional sensory molecule in bladder urothelium: Stretch-independent, tissue-specific actions and pathological implications. FASEB J. 34, 263–286 (2020).

49. K. Shibasaki, TRPV4 activation by thermal and mechanical stimuli in disease progression. Lab. Investig. 100, 218–223 (2020).

50. T. L. Toft-Bertelsen, et al., Lysophosphatidic acid as a CSF lipid in posthemorrhagic hydrocephalus that drives CSF accumulation via TRPV4-induced hyperactivation of NKCC1. Fluids Barriers CNS 19 (2022).

51. L. Lapajne, et al., TRPV4: Cell type-specific activation, regulation and function in the vertebrate eye. Curr. Top. Membr. 89, 189–219 (2022).

52. M. Lakk, et al., Membrane cholesterol regulates TRPV4 function, cytoskeletal expression, and the cellular response to tension. J. Lipid Res. 62 (2021).

53. T. Uchida, et al., TRPV4 is activated by mechanical stimulation to induce prostaglandins release in trabecular meshwork, lowering intraocular pressure. PLoS One 16 (2021).

54. L. Jing, K. Liu, F. Wang, Y. Su, Role of mechanically-sensitive cation channels Piezo1 and TRPV4 in trabecular meshwork cell mechanotransduction. Hum. Cell 37, 394–407 (2024).

55. S. N. Redmon, et al., TRPV4 subserves physiological and pathological elevations in intraocular pressure. Res. Sq. [Preprint] (2024). 10.21203/rs.3.rs-4714050/v1.

56. A. Nettesheim, M. S. Shim, J. Hirt, P. B. Liton, Transcriptome analysis reveals autophagy as regulator of TGFβ/Smad-induced fibrogenesis in trabecular meshwork cells. Sci. Rep. 9 (2019).

57. H. Li, J. L. Henty-Ridilla, A. M. Bernstein, P. S. Ganapathy, S. Herberg, TGFβ2 Regulates Human Trabecular Meshwork Cell Contractility via ERK and ROCK Pathways with Distinct Signaling Crosstalk Dependent on the Culture Substrate. Curr. Eye Res. 47, 1165–1178 (2022).

58. X. Yan, X. Xiong, Y. G. Chen, Feedback regulation of TGF-β signaling. Acta Biochim. Biophys. Sin. (Shanghai). 50, 37–50 (2018).

59. T. Carreon, E. van der Merwe, R. L. Fellman, M. Johnstone, S. K. Bhattacharya, Aqueous outflow - A continuum from trabecular meshwork to episcleral veins. Prog. Retin. Eye Res. 57, 108–133 (2017).

60. O. Yarishkin, et al., TREK-1 channels regulate pressure sensitivity and calcium signaling in trabecular meshwork cells. J. Gen. Physiol. 150, 1660–1675 (2018).

61. M. Honjo, et al., Effects of protein kinase inhibitor, HA1077, on intraocular pressure and outflow facility in rabbit eyes. Arch. Ophthalmol. 119, 1171–1178 (2001).

62. P. V. Rao, P. P. Pattabiraman, C. Kopczynski, Role of the Rho GTPase/Rho kinase signaling pathway in pathogenesis and treatment of glaucoma: Bench to bedside research. Exp. Eye Res. 158, 23–32 (2017).

63. C. R. Ethier, A. T. Read, D. W. H. Chan, Effects of latrunculin-B on outflow facility and trabecular meshwork structure in human eyes. Investig. Ophthalmol. Vis. Sci. 47, 1991–1998 (2006).

64. W. Liedtke, J. M. Friedman, Abnormal osmotic regulation in trpv4-/- mice. Proc. Natl. Acad. Sci. U. S. A. 100, 13698–13703 (2003).

65. D. A. Ryskamp, et al., The polymodal ion channel transient receptor potential vanilloid 4 modulates calcium flux, spiking rate, and apoptosis of mouse retinal ganglion cells. J. Neurosci. 31, 7089–7101 (2011).

66. O. Yarishkin, T. T. T. Phuong, M. Lakk, D. Križaj, TRPV4 does not regulate the distal retinal light response. Adv. Exp. Med. Biol. 1074, 553–560 (2018).

67. K. Ikegami, S. Masubuchi, Suppression of trabecular meshwork phagocytosis by norepinephrine is associated with nocturnal increase in intraocular pressure in mice. *Commun*. Biol. 5 (2022).

68. S. O. Rahaman, et al., TRPV4 mediates myofibroblast differentiation and pulmonary fibrosis in mice. J. Clin. Invest. 124, 5225–5238 (2014).

69. V. P. Willard, et al., Transient receptor potential vanilloid 4 as a regulator of induced pluripotent stem cell chondrogenesis. Stem Cells 39, 1447–1456 (2021).

70. N. A. Sharif, Identifying new drugs and targets to treat rapidly elevated intraocular pressure for angle closure and secondary glaucomas to curb visual impairment and prevent blindness. Exp. Eye Res. 232 (2023).

71. R. N. Weinreb, C. B. Toris, B. T. Gabelt, J. D. Lindsey, P. L. Kaufman, Effects of prostaglandins on the aqueous humor outflow pathways. Surv. Ophthalmol. 47 (2002).

72. M. C. Leske, et al., Factors for glaucoma progression and the effect of treatment: The early manifest glaucoma trial. Arch. Ophthalmol. 121, 48–56 (2003).

73. R. Fuchshofer, M. Birke, U. Welge-Lussen, D. Kook, E. Lütjen-Drecoll, Transforming growth factor-β2 modulated extracellular matrix component expression in cultured human optic nerve head astrocytes. Investig. Ophthalmol. Vis. Sci. 46, 568–578 (2005).

74. W. E. Medina-Ortiz, R. Belmares, S. Neubauer, R. J. Wordinger, A. F. Clark, Cellular fibronectin expression in human trabecular meshwork and induction by transforming growth factor-β2. Investig. Ophthalmol. Vis. Sci. 54, 6779–6788 (2013).

75. B. Callaghan, et al., Genome-wide transcriptome profiling of human trabecular meshwork cells treated with TGF-β2. Sci. Rep. 12 (2022).

76. M. Inoue-Mochita, et al., P38 MAP kinase inhibitor suppresses transforming growth factor-β2-induced type 1 collagen production in trabecular meshwork cells. PLoS One 10 (2015).

77. M. H. Kang, D. J. Oh, J. heon Kang, D. J. Rhee, Regulation of SPARC by transforming growth factor β2 in human trabecular meshwork. Investig. Ophthalmol. Vis. Sci. 54, 2523–2532 (2013).

78. G. Patel, et al., Molecular taxonomy of human ocular outflow tissues defined by single-cell transcriptomics. Proc. Natl. Acad. Sci. U. S. A. 117, 12856–12867 (2020).

79. E. Reina-Torres, et al., The vital role for nitric oxide in intraocular pressure homeostasis. Prog. Retin. Eye Res. 83 (2021).

80. T. van Zyl, et al., Cell atlas of aqueous humor outflow pathways in eyes of humans and four model species provides insight into glaucoma pathogenesis. Proc. Natl. Acad. Sci. U. S. A. 117, 10339–10349 (2020).

81. W. Zhu, et al., The role of Piezo1 in conventional aqueous humor outflow dynamics. iScience 24 (2021).

82. P. P. Pattabiraman, et al., Rhoa gtpase-induced ocular hypertension in a rodent model is associated with increased fibrogenic activity in the trabecular meshwork. Am. J. Pathol. 185, 496–512 (2015).

83. M. Zhang, R. Maddala, P. V. Rao, Novel molecular insights into RhoA GTPase-induced resistance to aqueous humor outflow through the trabecular meshwork. Am. J. Physiol. - Cell Physiol. 295 (2008).

84. A. D. Güler, et al., Heat-evoked activation of the ion channel, TRPV4. J. Neurosci. 22, 6408–6414 (2002).

85. R. Nishimoto, et al., Thermosensitive TRPV4 channels mediate temperature-dependent microglia movement. Proc. Natl. Acad. Sci. U. S. A. 118 (2021).

86. E. Abad, et al., Activation of store-operated Ca2+ channels in trabecular meshwork cells. Investig. Ophthalmol. Vis. Sci. 49, 677–686 (2008).

87. M. Feger, et al., The production of fibroblast growth factor 23 is controlled by TGF-andbeta. Sci. Rep. 7 (2017).

88. S. Sharma, et al., TRPV4 ion channel is a novel regulator of dermal myofibroblast differentiation. Am. J. Physiol. - Cell Physiol. 312, C562–C572 (2017).

89. R. K. Adapala, et al., TRPV4 channels mediate cardiac fibroblast differentiation by integrating mechanical and soluble signals. J. Mol. Cell. Cardiol. 54, 45–52 (2013).

90. C. J. O’Conor, H. A. Leddy, H. C. Benefield, W. B. Liedtke, F. Guilak, TRPV4-mediated mechanotransduction regulates the metabolic response of chondrocytes to dynamic loading. Proc. Natl. Acad. Sci. U. S. A. 111, 1316–1321 (2014).

91. Y. Songa, et al., TRPV4 channel inhibits TGF-β1-induced proliferation of hepatic stellate cells. PLoS One 9 (2014).

92. Y. Wu, J. Qi, C. Wu, W. Rong, Emerging roles of the TRPV4 channel in bladder physiology and dysfunction. J. Physiol. 599, 39–47 (2021).

93. J. L. Jones, et al., TRPV4 increases cardiomyocyte calcium cycling and contractility yet contributes to damage in the aged heart following hypoosmotic stress. Cardiovasc. Res. 115, 46–56 (2019).

94. S. Chaigne, S. Barbeau, T. Ducret, R. Guinamard, D. Benoist, Pathophysiological Roles of the TRPV4 Channel in the Heart. Cells 12 (2023).

95. X. Wen, et al., Aortic smooth muscle TRPV4 channels regulate vasoconstriction in high salt-induced hypertension. Hypertens. Res. 46, 2356–2367 (2023).

96. Y. L. Chen, et al., Novel Smooth Muscle Ca2+-Signaling Nanodomains in Blood Pressure Regulation. Circulation 146, 548–564 (2022).

97. Y. Zhu, et al., Vascular Smooth Muscle TRPV4 (Transient Receptor Potential Vanilloid Family Member 4) Channels Regulate Vasoconstriction and Blood Pressure in Obesity. Hypertension 80, 757–770 (2023).

98. X. Zhao, K. E. Ramsey, D. A. Stephan, P. Russell, Gene and protein expression changes in human trabecular meshwork cells treated with transforming growth factor-β. Investig. Ophthalmol. Vis. Sci. 45, 4023–4034 (2004).

99. A. Zhavoronkov, et al., Pro-fibrotic pathway activation in trabecular meshwork and lamina cribrosa is the main driving force of glaucoma. Cell Cycle 15, 1643–1652 (2016).

100. J. A. Last, et al., Elastic modulus determination of normal and glaucomatous human trabecular meshwork. Investig. Ophthalmol. Vis. Sci. 52, 2147–2152 (2011).

101. U. Raychaudhuri, J. C. Millar, A. F. Clark, Tissue transglutaminase elevates intraocular pressure in mice. Investig. Ophthalmol. Vis. Sci. 58, 6197–6211 (2017).

102. A. T. Read, D. W. H. Chan, C. R. Ethier, Actin structure in the outflow tract of normal and glaucomatous eyes. Exp. Eye Res. 84, 214–226 (2007).

103. K. Wang, A. T. Read, T. Sulchek, C. R. Ethier, Trabecular meshwork stiffness in glaucoma. Exp. Eye Res. 158, 3–12 (2017).

104. R. Fuchshofer, D. A. Stephan, P. Russell, E. R. Tamm, Gene expression profiling of TGFβ2- and/or BMP7-treated trabecular meshwork cells: Identification of Smad7 as a critical inhibitor of TGF-β2 signaling. Exp. Eye Res. 88, 1020–1032 (2009).

105. J. Gottanka, D. Chan, M. Eichhorn, E. Lütjen-Drecoll, C. R. Ethier, Effects of TGF-β2 in Perfused Human Eyes. Investig. Ophthalmol. Vis. Sci. 45, 153–158 (2004).

106. L. M. Grove, et al., Translocation of TRPV4-PI3Kγ complexes to the plasma membrane drives myofibroblast transdifferentiation. Sci. Signal. 12 (2019).

107. A. K. Shukla, et al., Arresting a transient receptor potential (TRP) channel: β-arrestin 1 mediates ubiquitination and functional down-regulation of TRPV4. J. Biol. Chem. 285, 30115–30125 (2010).

108. Y. Matsumura, et al., The prophylactic effects of a traditional Japanese medicine, goshajinkigan, on paclitaxel-induced peripheral neuropathy and its mechanism of action. Mol. Pain 10 (2014).

109. Y. Zhang, et al., A transient receptor potential vanilloid 4 contributes to mechanical allodynia following chronic compression of dorsal root ganglion in rats. Neurosci. Lett. 432, 222–227 (2008).

110. N. Alessandri-Haber, O. A. Dina, E. K. Joseph, D. Reichling, J. D. Levine, A transient receptor potential vanilloid 4-dependent mechanism of hyperalgesia is engaged by concerted action of inflammatory mediators. J. Neurosci. 26, 3864–3874 (2006).

111. A. Maqboul, B. Elsadek, Expression profiles of TRPV1, TRPV4, TLR4 and ERK1/2 in the dorsal root ganglionic neurons of a cancer-induced neuropathy rat model. PeerJ 2018 (2018).

112. Y. Y. Cui, et al., Expression and functional characterization of transient receptor potential vanilloid 4 in the dorsal root ganglion and spinal cord of diabetic rats with mechanical allodynia. Brain Res. Bull. 162, 30–39 (2020).

113. T. Borrás, L. K. Buie, M. G. Spiga, J. Carabana, Prevention of nocturnal elevation of intraocular pressure by gene transfer of dominant-negative RhoA in rats. JAMA Ophthalmol. 133, 182–190 (2015).

114. B. C. Samuels, et al., Dorsomedial/perifornical hypothalamic stimulation increases intraocular pressure, intracranial pressure, and the translaminar pressure gradient. Investig. Ophthalmol. Vis. Sci. 53, 7328–7335 (2012).

115. K. Ikegami, Circadian rhythm of intraocular pressure. J. Physiol. Sci. 74, 14 (2024).

116. K. Z. Shen, R. A. North, A. Surprenant, Potassium channels opened by noradrenaline and other transmitters in excised membrane patches of guinea-pig submucosal neurones. J. Physiol. 445, 581–599 (1992).

117. E. A. Steinberg, K. A. Wafford, S. G. Brickley, N. P. Franks, W. Wisden, The role of K2P channels in anaesthesia and sleep. Pflugers Arch. Eur. J. Physiol. 467, 907–916 (2015).

118. B. D. Matthews, et al., Ultra-rapid activation of TRPV4 ion channels by mechanical forces applied to cell surface β1 integrins. Integr. Biol. 2, 435–442 (2010).

119. C. Goswami, J. Kuhn, P. A. Heppenstall, T. Hucho, Importance of non-selective cation channel TRPV4 interaction with cytoskeleton and their reciprocal regulations in cultured cells. PLoS One 5 (2010).

120. A. O. Jo, et al., Differential volume regulation and calcium signaling in two ciliary body cell types is subserved by TRPV4 channels. Proc. Natl. Acad. Sci. U. S. A. 113, 3885–3890 (2016).

121. A. Boussommier-Calleja, et al., Pharmacologic manipulation of conventional outflow facility in ex vivo mouse eyes. Investig. Ophthalmol. Vis. Sci. 53, 5838–5845 (2012).

122. D. R. Overby, et al., The structure of the trabecular meshwork, its connections to the ciliary muscle, and the effect of pilocarpine on outflow facility in mice. Investig. Ophthalmol. Vis. Sci. 55, 3727–3736 (2014).

123. K. E. Keller, et al., Consensus recommendations for trabecular meshwork cell isolation, characterization and culture. Exp. Eye Res. 171, 164–173 (2018).

124. T. Bagué, et al., Effects of Netarsudil-Family Rho Kinase Inhibitors on Human Trabecular Meshwork Cell Contractility and Actin Remodeling Using a Bioengineered ECM Hydrogel. Front. Ophthalmol. 2 (2022).

125. W. D. Stamer, R. E. B. Seftor, S. K. Williams, H. A. M. Samaha, R. W. Snyder, Isolation and culture of human trabecular meshwork cells by extracellular matrix digestion. Curr. Eye Res. 14, 611–617 (1995).

126. T. T. T. Phuong, et al., Calcium influx through TRPV4 channels modulates the adherens contacts between retinal microvascular endothelial cells. J. Physiol. 595, 6869–6885 (2017).

127. J. Schindelin, et al., Fiji: An open-source platform for biological-image analysis. Nat. Methods 9, 676–682 (2012).

